# The genome of the contractile demosponge *Tethya wilhelma* and the evolution of metazoan neural signalling pathways

**DOI:** 10.1101/120998

**Authors:** Warren R. Francis, Michael Eitel, Sergio Vargas, Marcin Adamski, Steven H.D. Haddock, Stefan Krebs, Helmut Blum, Dirk Erpenbeck, Gert Wörheide

## Abstract

Porifera are a diverse animal phylum with species performing important ecological roles in aquatic ecosystems, and have become models for multicellularity and early-animal evolution. Demosponges form the largest class in sponges, but previous studies have relied on the only draft demosponge genome of *Amphimedon queenslandica*. Here we present the 125-megabase draft genome of a contractile laboratory demosponge *Tethya wilhelma*, sequenced to almost 150x coverage. We explore the genetic repertoire of transporters, receptors, and neurotransmitter metabolism across early-branching metazoans in the context of the evolution of these gene families. Presence of many genes is highly variable across animal groups, with many gene family expansions and losses. Three sponge classes show lineage-specific expansions of GABA-B receptors, far exceeding the gene number in vertebrates, while ctenophores appear to have secondarily lost most genes in the GABA pathway. Both GABA and glutamate receptors show lineage-specific domain rearrangements, making it difficult to trace the evolution of these gene families. Gene sets in the examined taxa suggest that nervous systems evolved independently at least twice and either changed function or were lost in sponges. Changes in gene content are consistent with the view that ctenophores and sponges are the earliest-branching metazoan lineages and provide additional support for the proposed clade of Placozoa/Cnidaria/Bilateria.

## Introduction

The presence of neurons is a defining character of animals, and is symbolic of their alleged superiority over all other life on earth. Nonetheless, the four non-bilaterian phyla, Porifera, Placozoa, Ctenophora and Cnidaria, are most different from other animals in their sensory systems and are often mistakenly referred to as “lower” animals in common parlance, despite the fact that, like bilaterians, non-bilaterians exist at the tips of the tree of life. Indeed, animals such as corals and sponges appear immobile or often unresponsive, challenging early theorists in their ideas of what is and is not an animal. Yet we now know that representatives from all four non-bilaterian phyla demonstrate dynamic responses to outside stimuli.

Neural evolution has been discussed previously in the context of paleontology (reviewed in [Wray et al., 2015]) and metazoan phylogeny (reviewed in [Jékely et al., 2015]). Indeed, it has been suggested that many features of bilaterian neurons and nervous systems represent separate, parallel evolutionary events from a “simple” nervous system. A simple nervous system then must arise from proto-neurons [Schierwater et al., 2009], however it is unclear what that might look like.

Several qualities can be used to define neurons or proto-neurons [Leys, 2015, Nickel, 2010] such as synapses, electrical excitability, membrane potential, or secretory functions, though no single quality (and ultimately gene set) solely defines such cells as neurons. Two non-bilaterian groups, ctenophores and cnidarians, are thought to have true neurons. When considering the remaining two non-bilaterian phyla, sponges and placozoans, many components of neural cells are found without any neuron-like cells having been identified [Srivastava et al., 2010, Riesgo et al., 2014a, Leys, 2015], although synapse-like structures have been identified in placozoan fiber cells that show vesicles close to an osmophile contact [Grell and Benwitz, 1974].

Comparative analyses revealed a gradient of neural-like qualities indicating that “neuron-or-not” classifications are not straightforward. While ctenophores, cnidarians, and bilaterians have true neurons, structural and biochemical differences, [Moroz et al., 2014, Moroz, 2015] led to the proposition that neurons in ctenophores and cnidarians may not be homologous, but rather separate evolutionary outcomes from neural-like precursor cells. Potentially, in the case of independent evolutions, neurons are “easy” to evolve, since it involves co-expression of various pan-metazoan genetic modules in the same cell type. Alternatively, early rudimentary signaling systems may have been energetically costly and not especially useful in pre-Cambrian oceans, and in such cases, it may have been comparatively easy to lose such genes and with them neuronal-type cells.

Interpretation of neural evolution requires an accurate metazoan phylogeny, and the phylogenetic relationships of early-branching metazoans have been a topic of continued controversy. Some analyses support the traditional phylogenetic position of sponges as sister group to all other metazoans (“Porifera-sister”) [Philippe et al., 2009, Pick et al., 2010, Nosenko et al., 2013, Pisani et al., 2015, Simion et al., 2017] while others suggest that Ctenophora are the sister group to all other animals (“Ctenophora-sister”) [Dunn et al., 2008, Ryan et al., 2013, Whelan et al., 2015], and some analyses also recover the classical view, a Coelenterata clade uniting Cnidaria and Ctenophora [Philippe et al., 2009, Simion et al., 2017]. Importantly, phylogenomic analyses can be prone to systematic artifacts under some circumstances, depending on taxon sampling [Pick et al., 2010, Philippe et al., 2011], gene set [Nosenko et al., 2013], phylogenetic model [Pisani et al., 2015], or use of nucleotides instead of proteins [Jarvis et al., 2014]. Other methods based on presence or absence of the genes themselves have been proposed to provide a sequence-independent inference of phylogeny [Ryan et al., 2010, Ryan et al., 2013, Pisani et al., 2015], relying on the assumption that gene loss is a rare event. However, non-bilaterians have the additional problem that basic knowledge of many aspects of their biology is absent [Dunn et al.,2015], and so the biological context that may separate or unite groups is limited.

In the context of phylogeny, the branching order critically affects whether neurons evolved multiple times or were lost (see schematic in Figure 1). Given the gradient of neural-like qualities, the actual evolutionary scenario may be somewhere in between a simple gain-loss of neurons. While some previous studies have focused on neural evolution in ctenophores [Ryan et al., 2013, Alberstein et al., 2015, Li et al., 2015] or analysing the genomic data from *A. queenslandica* [Krishnan et al., 2014], these alone do not provide a comprehensive picture of all animals.

**Figure 1:**
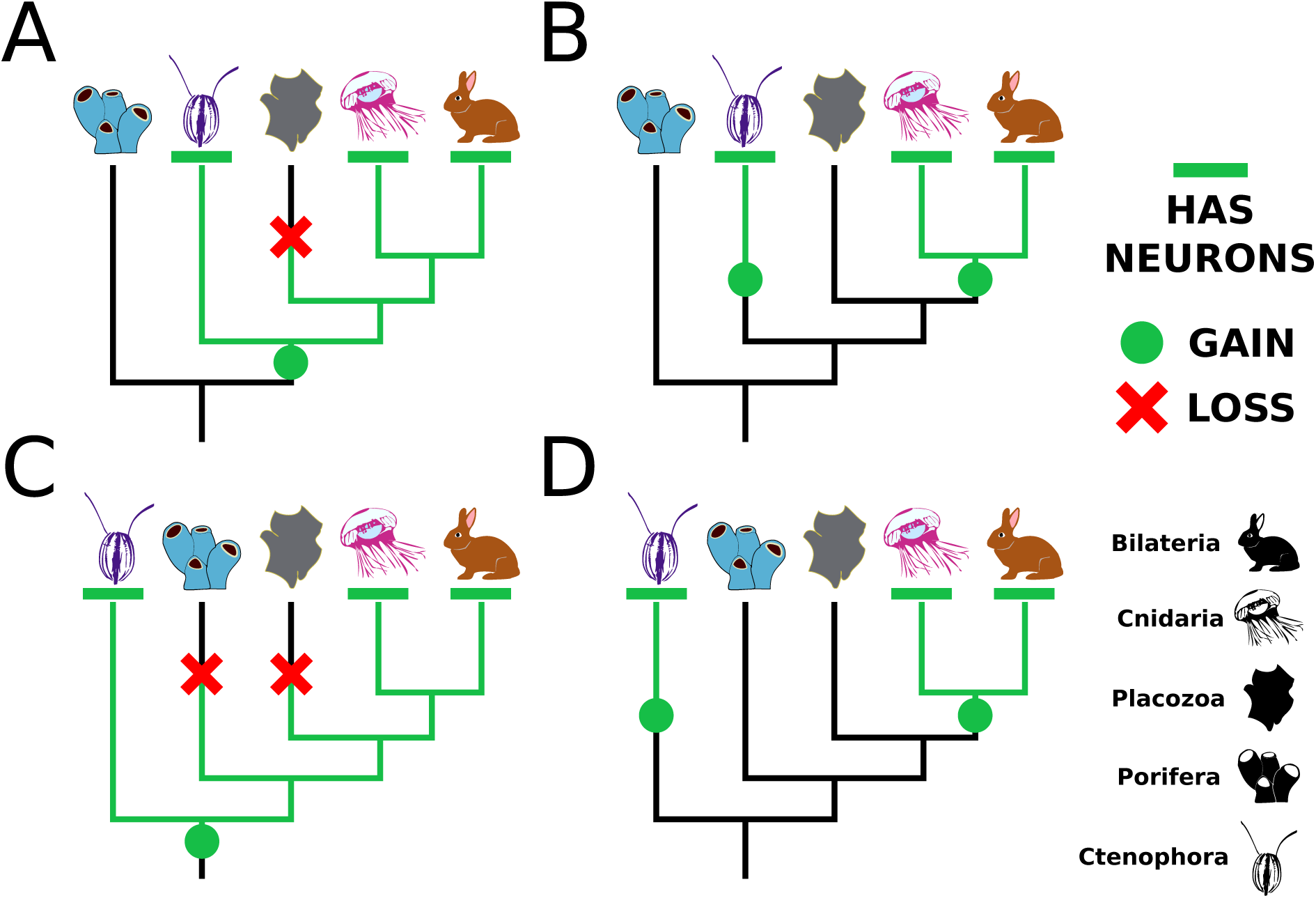
Schematic of neural evolution depending on metazoan phylogeny. The presence of neurons or neural-like cells in ctenophores, cnidarians, and bilaterians can be viewed differently depending on the phylogeny. Two different metazoan phylogenies based on recent multi-gene phylogenetic analyses are the source of the Porifera-sister (A,B) [Philippe et al., 2009, Pisani et al., 2015, Simion et al., 2017] and Ctenophora-sister (C,D) [Ryan et al., 2013, Whelan et al., 2015] scenarios. Neurons can either have evolved once requiring a secondary loss in sponges, placozoans, or both (A,C), or evolved twice, in ctenophores and in cnidarians/bilaterians (B,D).

Here we have sequenced the genome of the contractile laboratory demosponge *Tethya wilhelma* [Sara et al., 2001] and examined the protein repertoire in the context of genes mediating the contraction, and other neural-like functions. Many metabolic genes show unique expansions in different sponge clades, as well as other phyla, making it challenging to clearly assign functions based on similarity to human proteins. We consider these expansions in the context of phylogenetics, showing that even though sponges lack neurons, signaling pathways have still expanded. This gives support to the hypothesis that early neural-like cells have become neurons multiple times in the history of animals.

## Results

### Genome assembly and annotation

We generated a total of 61 gigabases of paired-end reads from a whole specimen of *T. wilhelma* (Figure 2) and all associated bacteria. Because of a close association with microbes, some contigs were expected to have derived from bacteria, as many reads have unexpectedly high GC content (Supplemental Figures 1-4). After assembly and filtration of bacterial contigs, the final assembly was 125Mb, similar to *A. queenslandica*, with a N50 value of 70kb (Supplemental Table 1). Gene annotation was done with a combination of a deeply-sequenced RNAseq library from an adult sponge and *ab initio* gene predictions. Because of high density of genes, extensive manual curation was often necessary to correct genes of the same strand that were erroneously merged. After correction and filtering of the *ab initio* predictions, we counted 37,416 predicted genes, comparable with the counts in *A. queenslandica* (40,122) [Fernandez-Valverde et al., 2015] and *S. ciliatum* (40,504) [Fortunato et al., 2014].

**Figure 2:**
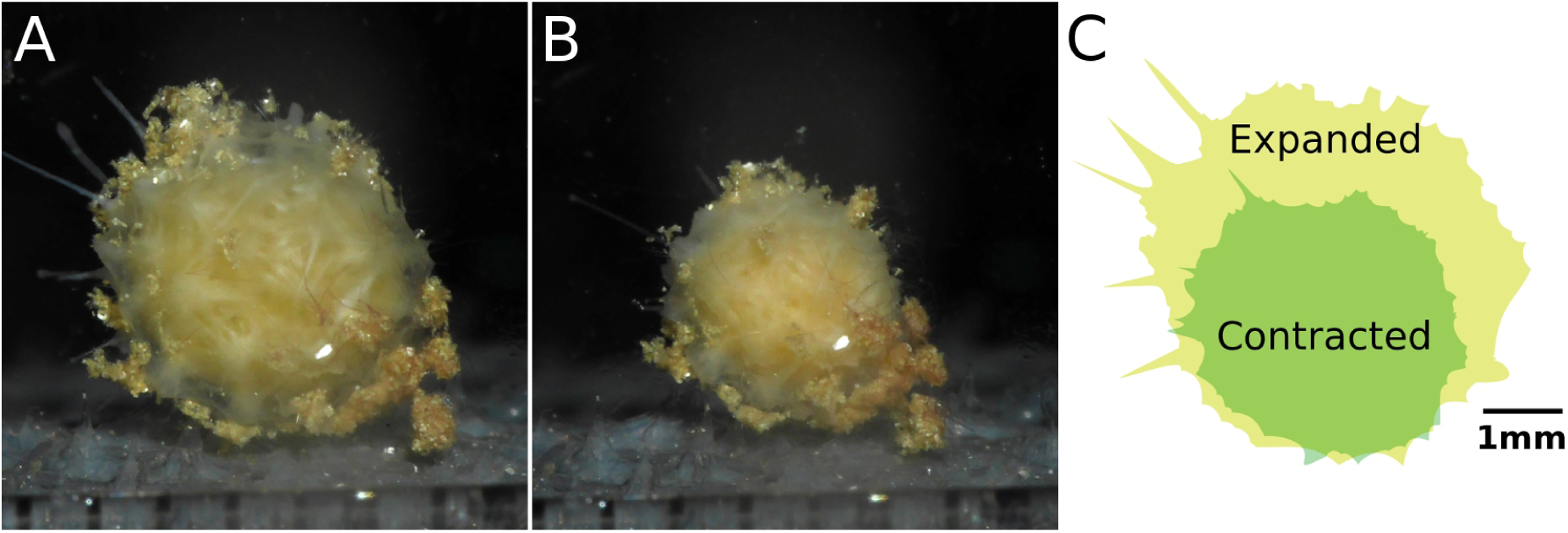
Contraction of a normal specimen. (A) Maximally expanded state of *Tethya wilhelma*. (B) The same specimen approximately an hour later in the most contracted state. (C) Cartoon view of most contracted versus expanded state. Scale bar applies to all images. Photos courtesy of Dan B. Mills.

General trends in splice variation were similar between *T. wilhelma* and *A. queenslandica* (Supplemental Tables 2 and 3), suggesting similar underlying biology or genome structure. One-to-one orthologs from *T. wilhelma* and *A. queenslandica* had relatively low identity (Supplemental Figure 5), with the average identity of 57.8%, showing a high genetic diversity within Porifera. The average identity is lower when compared to *S. ciliatum* (49.7%), *N. vectensis* (53.5%) and human (52.0%), which is not surprising given that *A. queens-landica* and *T. wilhelma* are both demosponges. Although both genomes are too fragmented to find syntenic chromosomal regions, ordered blocks of genes are still identifiable between *T. wilhelma* and *A. queenslandica* (Supplemental Figure 6), though not with *S. ciliatum*.

### Neurotransmitter metabolism across early-branching metazoans

Compared to most other metazoans, sponges have a limited set of behaviors (contraction, closure of osculum or choanocyte chambers to control flow), yet respond to many signaling molecules present in bilaterians [Ellwanger and Nickel, 2006, Ellwanger et al., 2007]. Some genes involved in vertebrate-like neurotransmitter metabolism have been found in sponges [Riesgo et al., 2014a, Krishnan and Schiöth, 2015]. although many display a sister-group relationship to homologs found in other animals and appear to have a complex evolutionary history with duplications in sponges and other non-bilaterian animals (Figure 3, Supplemental Figures 7-9), making the prediction of their functions difficult. Implicitly, presence of a gene usually means a one-to-one orthology relationship with a functionally annotated protein, probably a human protein. Since many of the non-bilaterian proteins in our set are many-to-many orthologs to human proteins with known functions, declaring presence or absence of any individual gene or genetic module is not correct in the strictest sense, as one-to-many or many-to-many orthologs are not the same gene. In such cases, it is not currently possible to computationally predict which, if any, of the sponge orthologs shares its function with a human protein.

**Figure 3:**
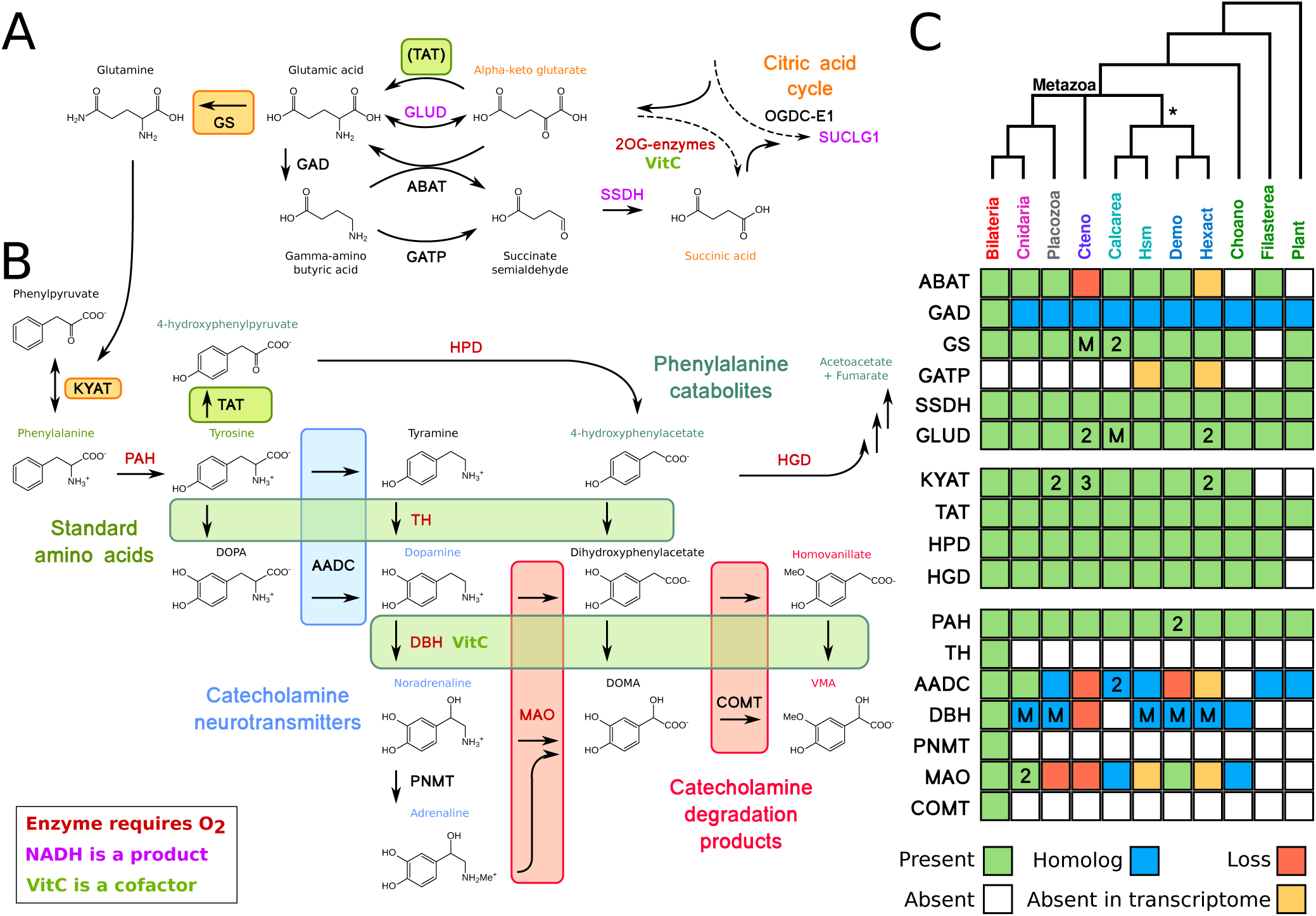
Neurotransmitter overview across metazoans. Summary schematic of neurotransmitter biosynthesis and degradation pathways across early-branching metazoans. Each of the four non-bilaterian groups is presumed to be monophyletic, although some individual trees of genes or gene families may display alternate topologies. Bold letters refer to the enzymes. Individual gene trees that display the orthology of the clades are found in the supplemental information. Arrows are shown in one direction though many reactions can be reversible. (A) Glutamate and GABA metabolism from the citric acid cycle through the “GABA shunt” pathway. Because ABAT is absent in ctenophores, GLUD and TAT are potentially alternatives to convert *α*-ketoglutarate to glutamate. GATP can convert GABA to succinate semialdehyde, but this enzyme was only found in some demosponges and plants. (B) Monoamine metabolism, excluding tryptophan. (C) Table of presence-absence for genes in parts A and B. Presence (green) refers to a 1-to-1 ortholog where orthology is clear from the tree position. Homolog (blue) refers to a sister group position in trees before duplications with different or unknown functions. Secondary loss (red) refers to the gene missing in the clade, but homologs are found in non-metazoan phyla. Numbers inside the boxes indicate copy number specific to that group, M refers to multiple duplications within the group where the copy number is variable among species. Abbreviations for clades are as follows: Cteno, ctenophores; Hsm, homoscleromorphs; Demo, demosponges; Hexact, hexactinellids; Choano, choanoflagellates.

For instance, biosynthesis of monoamine neurotransmitters (dopamine, serotonin, etc.) requires two enzymes, tryptophan hydroxylase and tyrosine hydroxylase. These two enzymes appear to have arisen in bilaterians from duplications of an ancestral phenylalanine hydroxylase [Cao et al., 2010], though evidence is lacking as to whether this ancestral protein had multiple functions that specialized after duplication (sub-functionalization) or developed new functions (neofunctionalization) post-duplication. The absence of these proteins in non-bilaterians seems to be ancestral; in other words, they had not evolved yet when these groups split and diversified.

Among other non-bilaterians, some monoamine neurotransmitters are found in cnidarians [Carlberg and Rosengren, 1985], but are mostly absent in ctenophores (or at detection limit) [Moroz et al., 2014]. Indeed, previous studies were unable to find homologs of DOPA decarboxylase (AADC, Supplemental Figure 8), dopamine *β*-hydroxylase (DBH, Supplemental Figure 7), monoamine oxidase (MAO, Supplemental Figure 9), or tyrosine hydroxylase (TH) in the genome of the ctenophore *M. leidyi* or any available ctenophore transcriptome, and it was suggested that some of these proteins were absent in sponges as well (see Supplementary Tables 17 and 19 in [Ryan et al., 2013]). However, we found orthologs of MAO and homologs of AADC and DBH in several sponges, though it is unclear if they perform the same function as the human proteins. Additionally, homologs of four enzymes, AADC, MAO, DBH, and ABAT, are present in single-celled eukaryotes but not ctenophores, implying a secondary loss of these protein families in this phylum.

#### GABA receptors

The neurotransmitter gamma-amino butyric acid (GABA) has been shown to affect contraction in *T. wilhelma* [Ellwanger et al., 2007] and the freshwater sponge *E. muelleri* [Elliott and Leys, 2010]. The genome of *T. wilhelma* contains metabotropic receptors (GABA-B, mGABARs), but not ionotropic GABA receptors (GABA-A, iGABARs). While humans have two mGABARs and the ctenophore *M. leidyi* has only one, the *T. wilhelma* genome has nine. Sponges appear to have undergone a large expansion of this protein family (Figure 4), similar to the expansion of glutamate-binding GPCRs previously observed in sponges [Krishnan et al., 2014]. Based on the structure of the binding pocket of human GABAR-B1 [Geng et al., 2013], many differences are observed across the mGABAR protein family, even showing that many residues involved in coordination of GABA are not conserved between the two human proteins or all other animals (Supplemental Figure 11). Contrary to previous reports [Ramoino et al., 2010], we were unable to find normal mGABARs in the two calcareous sponges *S. ciliatum* and *L. complicata*. Instead, in these two species, the best BLAST hits from human GABA-B receptors (the putative mGABARs) had the best reciprocal hits to Insulin-like growth-factor receptors. Structurally, this was due to the normal seven-transmembrane domain being swapped with a C-terminal protein kinase domain (Figure 5), meaning these are not true metabotropic GABA receptors. Similarly, in the filasterean *Capsaspora owczarzaki*, the N-terminal ligand binding domain is also exchanged with other domains, suggesting as well that these are not true metabotropic GABA receptors.

**Figure 4:**
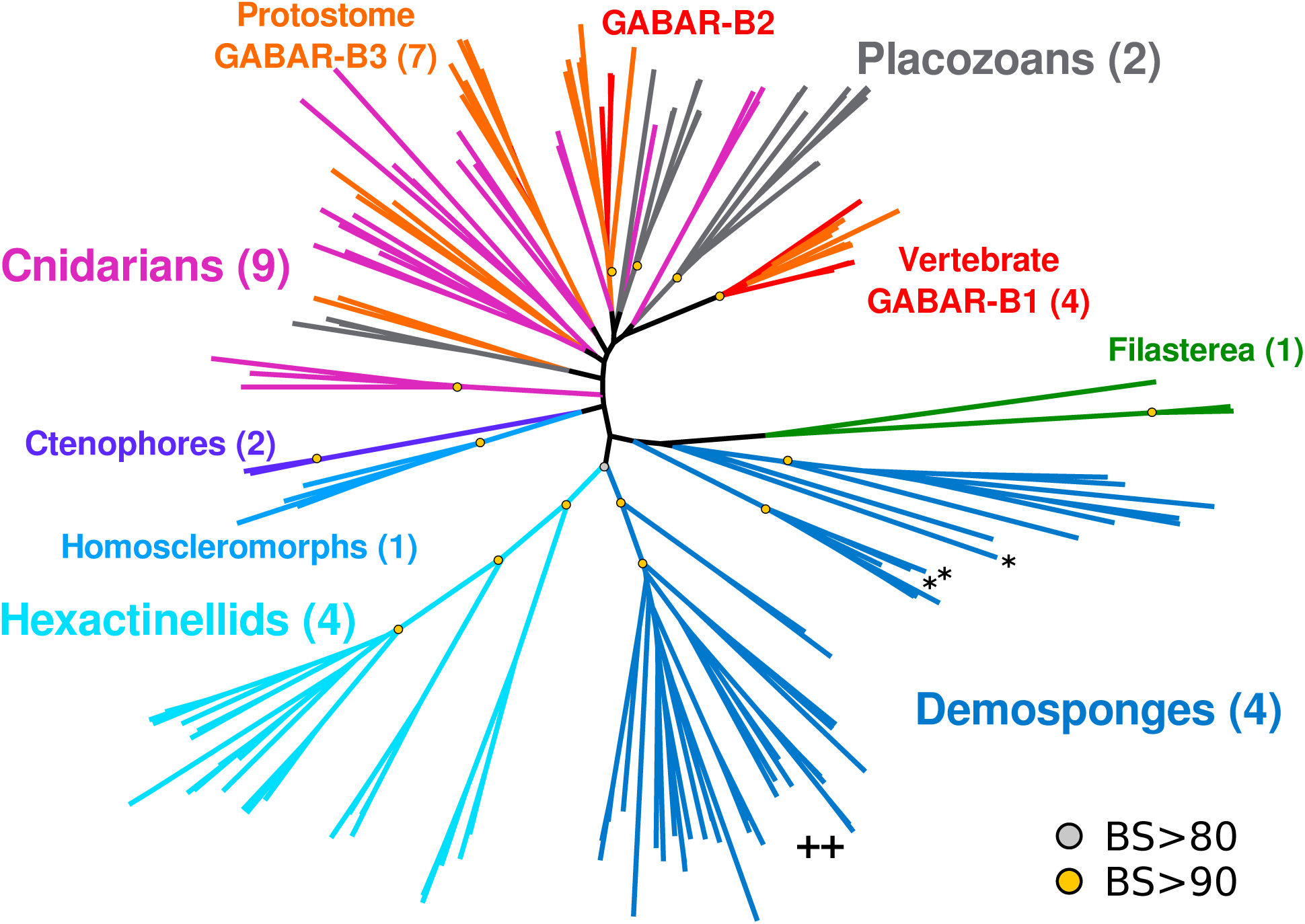
Metabotropic GABA receptors (GABA-B type) across metazoans. Protein tree generated with RAxML. Numbers in parentheses indicate the number of species from that group, so the 46 demosponge mGABARs come from 4 species. Key bootstrap values are summarized as yellow or gray dots, for values of 90 or more, or 80 or more, respectively. Single star indicates sequences annotated as mGABARs in [Krishnan et al., 2014], double plus indicates the clade annotated as “sponge specific expansion” in [Krishnan et al., 2014]. For complete version with protein names and all bootstrap values, see Supplemental Figure 10

**Figure 5:**
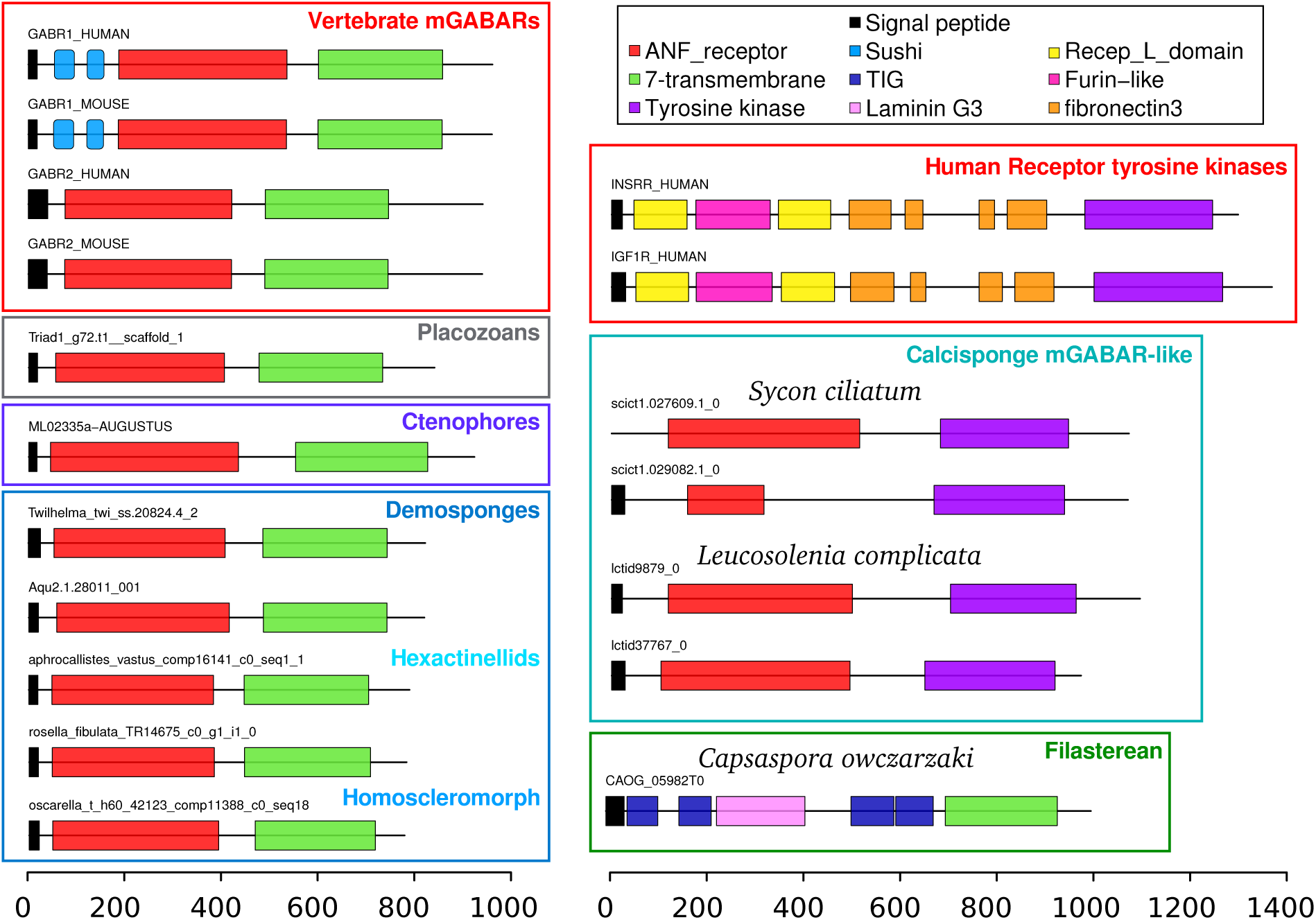
Domain organization of GABA-B type receptors across metazoans. Scale bar displays number of amino acids. Top reciprocal BLAST hits to human for putative mGABARs in calcareous sponges are INSRR and IGF1R, due to high-scoring hits to the tyrosine kinase domain. All mGABARs from demosponges, glass sponges, and the homoscleromorph *Oscarella carmela* share the 7-transmembrane domain (green) with mGABARs from other animals, while calcarean proteins have the same ligand-binding domain (red) but instead have a protein tyrosine kinase domain (purple) at the C-terminus, similar to growth-factor receptors. The filasterean *Capsaspora owczarzaki* has alternate domains at the N-terminus.

#### Glutamate receptors

Glutamate is of particular interest as it is a key metabolic intermediate and the main excitatory neurotransmitter in animal nervous systems, acting on two types of receptors: the metabotropic glutamate receptors (mGluRs) and the ionotropic ones (iGluRs). Some sponge species possess iGluRs, though these receptors were absent in the transcriptomes of several demosponges [Riesgo et al., 2014a]. We were unable to find iGluRs in the genome of *T. wilhelma*, in the genome and transcriptomes of any other demosponge, or in the genomes of two choanoflagellates (*M. brevicollis* and *S. rosetta*). The top BLAST hits in demosponges have a GPCR domain instead of the ion channel domain, indicating that these are not true iGluRs (Supplemental Figure 13). Because the domain structure in plants is the same as most animal iGluRs, the ligand binding domain was swapped out in demosponges.

The homoscleromorpha/calcarea clade appears to have an independent expansion of iGluRs (Supplemental Figure 12), though the normal ion transporter domain is switched with a SBP-bac-3 domain (PFAM domain PF00497) compared to all other iGluRs (Supplemental Figure 13). Additionally, ctenophores and placozoans appeared to have dramatic expansions of this protein family as well [Ryan et al., 2013, Moroz et al., 2014, Alberstein et al., 2015], suggesting that a small set of iGluRs was present in the common ancestor of eukaryotes and have diversified multiple times in both plants and animals, while other clades appear to have modified or lost these proteins.

#### Vesicular transporters

Secretory systems are a common feature of all eukaryotes, as most cells have endoplasmic reticulum to secrete proteins or make membrane proteins. Neurons secrete peptides (conceptually identical to any other protein) or small-molecule neurotransmitters in a paracrine fashion, specifically to other neural cells. Compared to peptides, small-molecule neurotransmitters need to be loaded into vesicles by dedicated transport proteins. Vesicular glutamate transporters (VGluTs, SLC17A6-8) are part of a superfamily of transporters [Sreedharan et al., 2010] that carry glutamate, aspartate, and nucleotides. The position of sponge proteins in the tree is inconsistent with a clear role in glutamate transport (Supplemental Figure 14), as several sponge clades and ctenophores occur as sister group to multiple duplications. Transporters in sponges, ctenophores, and choanoflagellates may well act upon glutamate or other amino acids, but this needs to be experimentally investigated.

Similar to glutamate, GABA is loaded into vesicles with the vesicular inhibitory amino acid transporter (VIAAT). Ctenophores, sponges, and placozoans lack one-to-one homologs of VIAAT (Supplemental Figure 15). Several other transporters are thought to transport GABA (ANTL or SLC6 class) and many other amino acids. SLC6-class transporters, which transport diverse amino acids, are found in all non-bilaterian groups, so the function of VIAAT may be redundant.

#### Glycine receptors

Glycine is known to affect the contraction of *T. wilhelma* [Ellwanger and Nickel, 2006]. Some ctenophore iGluRs have been shown to bind glycine [Alberstein et al., 2015] due to the substitution of serine for arginine (S687 in human GluN1), though this appears to be specific only to ctenophores, as essentially all other iGluRs have the conserved serine/threonine at this position. Because no ionotropic glycine receptors could be identified in the *T. wilhelma* genome (or any other sponges, ctenophores or placozoans), other proteins may be responsible for mediating this effect.

#### Mechanical receptors

Some sponges can contract in response to mechanical agitation, as reported for the demosponges *E. muelleri* [Elliott and Leys, 2007] and *T. wilhelma* [Nickel, 2010]. Several diverse protein families appear to be responsible for the sense of touch [Árnadóttir and Chalfie, 2010]. A subgroup of the TRP (transient receptor-potential) channels, TRP-N, thought to mediate mechanosensation was determined to be absent in sponges [Schuler et al., 2015], and we were unable to identify any in either *T. wilhelma* or *S. ciliatum*, although other TRP-class channels were found [Ludeman et al., 2014, Schuler et al., 2015]. Because the mechanosensory function of TRP channels may be redundant, we analysed for the presence of PIEZO, a 280kDa trimeric protein [Ge et al., 2015] involved in touch sensation in mammals [Coste et al., 2012]. Although two homologs were found in vertebrates, we found one copy in all other animals (Supplemental Figure 16) as well as fungi, plants and most other eukaryotic groups, suggesting an ancient and conserved function of this protein.

#### Voltage-gated channels

Voltage-gated ion channels are necessary for the propagation of electrical signals down axons and dendrites [Zakon, 2012, Moran et al., 2015], and have specificities for sodium, potassium, or calcium. Previous analyses were unable to clearly identify potassium or sodium channels in sponges [Liebeskind et al., 2011]; only one partial potassium channel was found in the transcriptome of the homoscleromorph *Corticium candelabrum* [Riesgo et al., 2014a, Li et al., 2015]. We were unable to find any voltage-gated sodium or potassium channels in the genome or transcriptome of any sponge. We then examine voltage-gated hydrogen channels (hvcn1), as these proteins have been found in a number of single-celled eukaryotes [Smith et al., 2011], and are extremely conserved. These channels were found in all sponge groups, although the high protein identity resulted in a poorly-resolved tree (Supplemental Figure 19).

Reports of action potentials in hexactinellids [Leys et al., 1999, Leys et al., 2007, Nickel, 2010] showed that sponge action potentials were inhibited by divalent cations [Leys et al., 1999], suggesting a role of calcium channels instead. Because voltage-gated sodium and calcium channels arose from a duplication event [Gur Barzilai et al., 2012], the ion selectivity may be variable within this protein family. Most sponges have only a single CaV-channel (Supplemental Figure 18) and several Hv-channels, and no voltage-gated channel of any kind was found in any glass sponge. However, all glass sponge sequences are from transcriptomes, therefore either the expression level of the true channels is low in glass sponges, or they have independently evolved another mechanism to propagate action potentials.

## Discussion

### Gene content variation of metazoa

Among the thousands of genes in the genome, we focused on genes that may be mediating contractile behavior in *T. wilhelma*, and the interactions of those genes within broader metabolic pathways. Many of the “housekeeping” genes in our study have lineage-specific duplications in at least one animal phylum. Considering the importance of “single-copy” proteins in phylogenetic analyses, as taxon sampling improves, it may be found that very few or no genes are single copy across most or all animal phyla. Many other genes that are critical for neural functioning in bilaterians have independent losses in other animal lineages (Figure 6).

**Figure 6:**
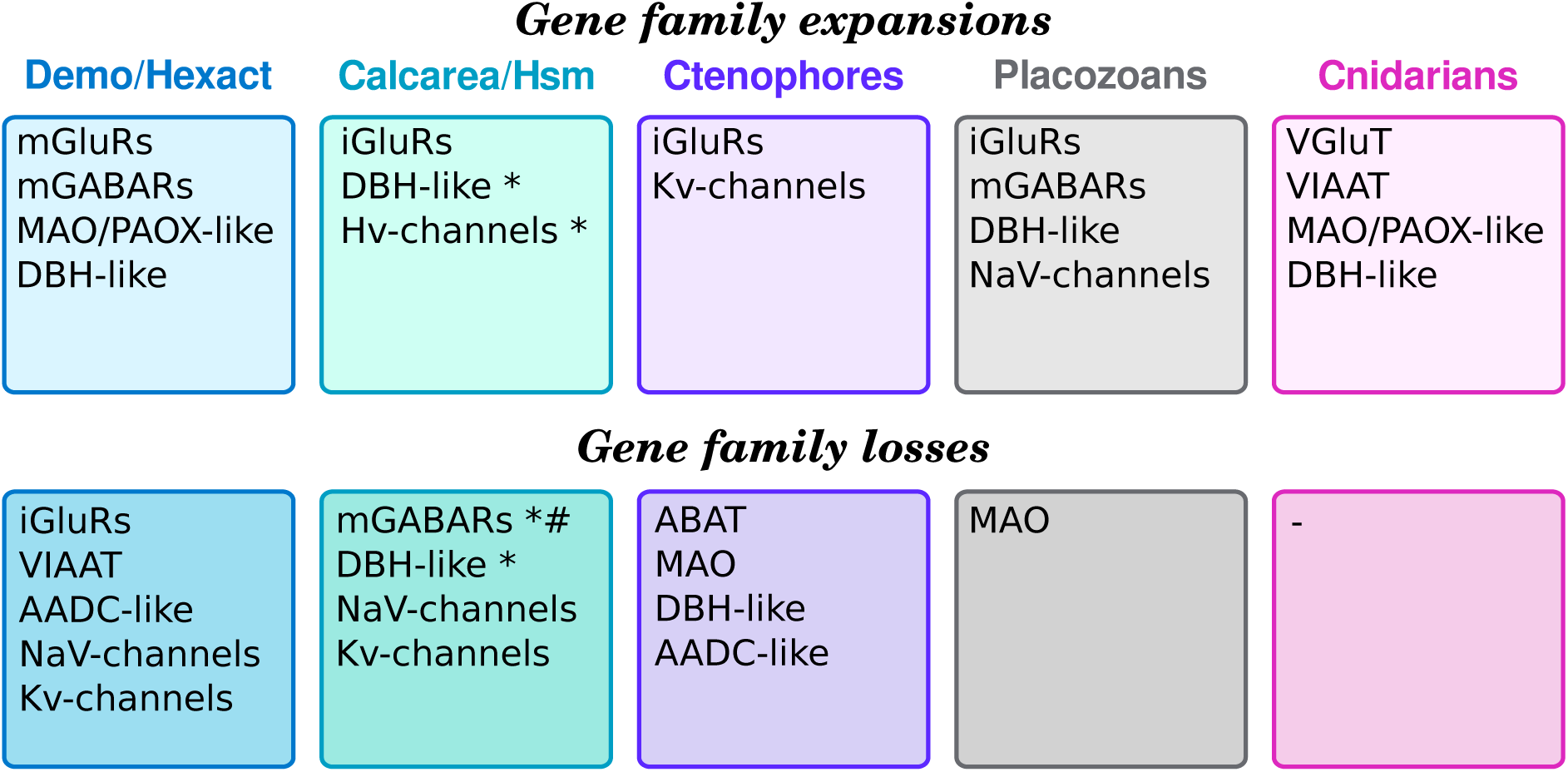
Summary of gene expansions and losses. Demo/Hexact refers to the clade of demosponges and glass sponges. Calcarea/Hsm refers to the unnamed clade of calcareous sponges and homoscleromorphs. The star indicates that expansion or loss was found in one class but not the other. Number sign indicates the domain rearrangement in Calcarea.

### Glutamate and GABA receptor evolution

There is stark contrast in the relative abundance of mGABARs and iGluRs in sponges and ctenophores. The relative dearth of mGABARs in ctenophores may reflect the apparently absence of amino-butyrate amino-transferase (ABAT) in ctenophores, suggesting that ctenophores use an alternate pathway to produce glutamate or metabolize GABA (Figure 3), rarely use GABA as a neurotransmitter, or simply are missing this pathway. Other aminotransferases such as GLUD or TAT may perform some of the exchange between *α*-ketoglutarate and glutamate, particularly as ctenophores have two copies of GLUD while most animals have only one. Ctenophores also have multiple (variable) copies of glutamate synthase and three copies of KYAT one of which may serve to balance glutamate metabolism in these animals.

There are two explanations for the diversity of mGABARs in sponges. Given the high variability of amino acids in the mGABAR binding pocket (Supplemental Figure 11), it is plausible that many of these receptors do not bind GABA at all, and have diversified for other ligands. There is precedent for this as it was shown that the independent expansion of ctenophore iGluRs also included several key mutations to the binding pocket which changed the ligand specificity of these proteins [Alberstein et al., 2015]. For the other hypothesis, all of the receptors could bind GABA, essentially mediating the same contraction signal, but their kinetics could differ and be influenced by factors such as, for instance, temperature. Because sponges are mostly immobile, they often can be subject to environment variation in terms of light, oxygen, and temperature. The possession of a set of proteins capable of triggering the same response (e.g. contraction) with varying daily or seasonal environmental conditions (e.g. temperature) would be beneficial and may explain the diverse set of receptors observed in sponges. Experimental characterization of these binding domains is necessary and may even show that a combination of these hypotheses explains the diversification of mGABARs in Porifera.

The apparent absence of true mGABARs in calcareous sponges (the genome of *S. ciliatum* and transcriptome of *L. complicata*) conflicts with a previous study that identified key proteins in the GABA pathway by immunostaining [Ramoino et al., 2010]. The best mGABARs BLAST hits found in the two calcareous sponges display a conserved ligand binding domain but the seven-transmembrane domain has been swapped with a tyrosine kinase domain (Figure 5). Structural similarity of the conserved N-terminal domain may result in a false-positive signal in studies using immunostaining with standard antibodies [Ramoino et al., 2010]. On the other hand, compared to ctenophores, which apparently lack ABAT, this enzyme was found in both of the calcareous sponges analyzed. Thus it would be surprising if these sponges had no capacity to create or respond to GABA. Since true vertebrate-like mGABARs are found in all other sponge classes, and our study could only examine two calcareous sponges, it could be that mGABAR presence is variable in this class. The genome of *S. ciliatum* contains 40 proteins annotated as mGluRs [Fortunato et al., 2014], so a third possibility is that even in the absence of true mGABARs, some of these proteins may have evolved affinity for GABA and mediate its signaling in calcareous sponges.

Although a putative iGluR was identified in the transcriptome of the demosponge *Ircinia fasciculata*, this sequence was only a fragment, so the glutamate affinity and domain structure could not be determined. As with the mGABARs, the domain structure is different between the sponge classes. Otherwise, it appears that only calcareous sponges and homoscleromorphs have NMDA/AMPA-like iGluRs. The presence of these proteins in plants and other single-celled eukaryotes suggests that at least iGluRs were present in the common ancestor of all eukaryotes, and their absence in demosponges is likely the product of secondary losses. In the context of contractions of *T. wilhelma*, the abundance of mGluRs and mGABARs could plausibly work in antagonistic ways via the action of different G-proteins making ionotropic channels not necessary for the modulation of this behavior.

### Variation in neurotransmitter metabolism

Many of the oxidative enzymes in the monoamine pathway require molecular oxygen, suggesting an important role of this molecule both the synthesis of the neurotransmitters (with PAH, TH, and DBH) and their inactivation (with MAO). Two catabolic pathways arise from tyrosine (Figure 3) and require oxygen at nearly all steps. It is unclear why intermediate products of one of these two pathways, the catecholamine pathway, became neurotransmitters and the other did not, particularly as hydroxyphenylpyruvate pathway is universally found in animals and catabolic intermediates are likely to be ubiquitous.

MAO was found in most animal groups, but we were unable to find any in placozoans or ctenophores. The topology of the MAO phylogenetic tree suggests a secondary loss of this protein in these phyla (Supplemental Figure 9). Related genes (PAOX, polyamine oxidase) were found in placozoans with several placozoan-specific duplications, and again, potentially one of these may catalyze the oxidation of aromatic amines. The analysis of these proteins also uncovered a clade including sponges, cnidarians, and lancelets, though the function of these proteins cannot be predicted based on homology searches. *In vitro* characterization of these enzymes may reveal the function to provide evidence as to how these could have been important for metabolism in early animals, and was subsequently replaced or lost in most other metazoan lineages.

Remarkably, the DBH group has independent expansions in three sponge classes as well as placozoans and cnidarians (Supplemental Figure 7). No DBH homologs were identified in calcareous sponges or in ctenophores. A putative homolog of this group was found in the choanoflagellate *M. brevicollis* but not in any other non-metazoan. The alignment and the phylogenetic position of the *M. brevicollis* protein suggest that it may be a member of the copper-binding oxygenase superfamily, rather than a true homolog of DBH (see Supplemental Alignment).

The presence of DBH-like and AADC-like enzymes in most animal groups suggests the possibility to make phenylethanolamines (like octopamine or noradrenaline) from tyrosine, and then subsequently inactivate them with MAO. All demosponges appear to lack AADC, and ctenophores appear to lack both of these enzymes calling into question a previous report of the detectability of monoamine neurotransmitters in ctenophores [Carlberg and Rosengren, 1985].

### Conserved properties of neurons

Neurons are defined by the presence of five key aspects: membrane potential, voltage-gated ion channels, secretory pathways, ligand-gated ion channels, and cell-cell junctions to form synapses. Voltage-gated channels, secretory systems, and ligand-gated ion channels are discussed above. Membrane potential is maintained in animal cells by sodium-potassium pumps (ATP-ases), which are a class of cation pumps exclusively found in animals [Stein, 1995, Sáez et al., 2009]. It is thought that such pumps are necessary because animals are the only multicellular group that lacks any kind of cell wall, thus careful control of ionic balance is necessary to resist osmotic stress [Stein, 1995]. For non-bilaterian animals, cell layers were in direct contact with water, so potentially all cells needed this protein to function normally. Therefore having neuron-like functionality is unlikely to rest upon the gain or loss of this gene. The last feature is the presence of cell-cell connections. Many proteins involved in synapse structure or neurotransmission are found in sponges, [Srivastava et al., 2010, Riesgo et al., 2014a, Moran et al., 2015, Leys, 2015] though it is not clear which genes are necessary for neural functioning, or may have evolved independently.

### Neural evolution and losses

Based on recent phylogenies, both Porifera-sister and Ctenophora-sister evolutionary scenarios require either at least one loss of neurons or two independent gains (Figure 1) of this cell-type. The only scenario that allows for a single evolution of neurons and no losses is the “Coelenterata” hypothesis (reviewed in [Jékely et al., 2015]), which joins cnidarians and ctenophores in a clade. However, many molecular datasets [Dunn et al., 2008, Ryan et al., 2013, Whelan et al., 2015, Pisani et al., 2015] and morphological evidence [Harbison, 1985] argue against this scenario (but also see [Philippe et al., 2009] and [Simion et al., 2017]). One other alternative is that placozoans have an unidentified neuron-like cell in a Porifera-sister context, which would therefore allow for a single origin of neural systems in animals and no losses.

What do the two different scenarios mean for evolution of neuronal cells? Considering the basic properties of neurons related to electrical signaling or secretory pathways, it had been shown before that many of the genes involved are universally found in animals. A single origin and multiple losses implies that the genetic toolkit necessary for all of these functions was present in the same single-celled organism or the same cell type (an hypothetical proto-neuron) of the last common ancestor of crown-group Metazoa, and either that cell type was lost or its functions were split up.

Sponges and ctenophores both appear to have lost several gene families (Figure 6), though ctenophores nonetheless have neural cells. Thus, the losses of the GABA or monoamine pathways are not critical for the functioning of neural cell types overall. However, voltage-gated potassium and sodium channels are thought to be essential for the propagation of electrical signals down axons and dendrites and have been found in all animal groups except sponges [Moran et al., 2015]. The NaV-channel tree shows a single origin of this protein family (Supplemental Figure 17), and presence of these channels in choanoflagellates suggests they were present in the common ancestor of all animals; the apparent absence in sponges therefore is probably a secondary loss. By comparison, ctenophores have a mostly-unique expansion of Kv-channels relative to the rest of metazoans [Li et al., 2015] and a duplication in NaV channels. Together with the loss of this protein family in sponges, the gene content argues for a combination of both multiple, independent gains and a loss of neural-type cells and their associated functions across animals.

Properties of the earliest metazoans are unknown, including life cycle or number of cell types, but it is most parsimonious that the first obligate multicellular animals did not have anything resembling a modern bilaterian nervous system [Wray et al., 2015]. Yet, the genomic evidence shows that these animals have cellular capacity to respond to environmental or paracrine signals, regulate the cell internal ion concentrations and respond to changes in their concentrations, and secrete small molecules that could serve as effectors in unconnected (but proximal) cells. Thus, the earliest animals likely had the capacity to develop nerve cells using the genetic toolkit they possessed, though the number of times this occurred is unclear. This capacity appears to have been lost in sponges with the loss of voltage-gated channels. As we were unable to find putative genes to mediate action potentials in glass sponges, either all of the four transcriptomes were incomplete or the unique action potentials of glass sponges may represent a third case of the evolution of neural-like functions in Metazoa.

## Methods

### Sequence data

Project overview can be found at spongebase.net. Reference data from the demosponge *Tethya wilhelma* are available at: https://bitbucket.org/molpalmuc/tethya_wilhelma-genome

Raw genomic reads for *T. wilhelma* are available on NCBI SRA under accession numbers SRR2163223 (genomic reads), SRR2296844 (mate pairs), SRR5369934 (DNA Moleculo), and SRR4255675 (RNAseq).

### Genome assembly

#### Processing and assembly

We generated 25Gb of 100bp paired-end Illumina reads of genomic DNA and 35Gb of 125bp Illumina gel-free mate-pair reads. Contigs were assembled with SOAPdenovo2 [Luo et al., 2012] using a kmer of 83bp. We also generated 436Mb of Moleculo synthetic long reads. Because both haplotypes are represented in the Moleculo reads, we merged the Moleculo reads using HaploMerger [Huang et al., 2012]. Contigs and merged Moleculo reads were then scaffolded using the gel-free mate-pairs with SSPACE [Boetzer et al., 2011] and BESST [Sahlin et al., 2014]. The first draft assembly had 7,947 contigs, totaling 145 megabases.

#### Removal of low-coverage contigs

To examine the completeness of the genome, we generated a plot of kmer coverage against GC percentage for the contigs (Supplemental Figure 1) using custom Python scripts (available at http://github.org/wrf/lavaLampPlot). This revealed 1,040 contigs with a coverage of zero that were carried over from the Moleculo reads and were not assembled (Supplemental Figure 2), accounting for 6 megabases. As these reads likely derived from bacterial contamination in the aquarium water, these contigs were removed, leaving 6,907 contigs totalling 138 megabases.

#### Separation of bacterial contigs

Additionally, the plot revealed many contigs with lower coverage (20x-90x) and high GC content (50-75%) suggesting the presence of bacteria (Supplemental Figure 3). Because many of these contigs were shorter than 10kb, separation of the bacterial contigs was done through several steps. We found 4,858 contigs with mapped RNAseq reads and GC content under 50%, as expected of metazoans. These contigs accounted for 88% of the sponge assembly, or 121 megabases. For the 2,014 contigs with no mapped RNAseq, we used blastn to search the contigs against the *A. queenslandica* scaffolds and all complete bacterial genomes from Genbank (5,242 sequences). Based on subtraction of bitscores, 62 contigs were identified as sponge and 565 were identified as bacterial. For the remaining 1,387 contigs, most of which were under 10kb, we repeated the search with tblastx against *A. queenslandica* scaffolds and the genomes of *Sinorhizobium medicae* and *Roseobacter litoralis*, which were the most similar complete genomes to the two bacterial 16S rRNAs identified in the contigs. After all sorting, 798 putative bacterial contigs accounted for 12.7 megabases and were separated to bring the total to 6,109 sponge contigs. Contigs for the two bacteria were binned by tetranucleotide frequency using MetaWatt [Strous et al., 2012] (Supplemental Figure 4).

#### Genome coverage and completeness

Coverage was estimated two ways: kmer frequency and read mapping. Kmers of 31bp were counted using the Jellyfish kmer counter [Marçais and Kingsford, 2011] and analyzed using custom Python and R scripts (“fastqdumps2histo.py” and “jellyfish_gc_coverage_blob_plot_v2.R”, available at http://github.org/wrf/lavaLampPlot). As expected, the kmer distribution showed two peaks, one for kmers at heterozygous positions and one for homozygous positions, whereupon the coverage peak was at 131-fold coverage for homozygous positions. Because of sequencing errors, this method often underestimates coverage, and so to confirm this estimate we then mapped all reads to the genome using Bowtie2 [Langmead and Salzberg, 2012]. The sum of mapped reads divided by the total length provided an estimated coverage of 159-fold physical coverage.

Of the original reads, 185 million (86.5%) mapped back to the assembled sponge contigs. Completeness for gene content was assessed with BUSCO [Simão et al., 2015], whereupon we found 728 (86%) complete genes and 42 (4.9%) predicted-incomplete genes. Overall, these data suggest that the genome assembly is adequate for downstream analyses.

### Genome annotation

#### Transcriptome versions

The transcriptome for *T. wilhelma* was assembled *de novo* using Trinity (release r20140717) [Grabherr et al., 2011, Haas et al., 2013]. Default parameters were used, except for strand specific assembly, *in silico* read normalization, and trimming (–SS_lib_type RF –normalize_reads –trimmomatic). This produced 127,012 transcripts with an average length of 913bp. Assembled transcripts were mapped to the genomic assembly using GMAP [Wu and Watanabe, 2005] to produce a GFF file of the transcript mapping. Of these, 114,744 transcripts were mapped 166,847 times, allowing for multiple mappings.

For the genome-guided transcriptome, strand-specific RNAseq reads were mapped against the genome build using Tophat2 v2.0.13 [Kim et al., 2013] using strand-specific mapping (option –library-type fr-firststrand) and otherwise default parameters. Mapped reads were then joined into transcripts using StringTie v1.0.2 [Pertea et al., 2015] with default parameters.

Additionally, *ab initio* gene models were predicted using AUGUSTUS [Stanke et al., 2008]. AUGUSTUS was trained on the webAugustus server [Hoff and Stanke, 2013] using the highest expression transcripts for each Trinity component and the assembled contigs. This identified 27,551 putative genes. The majority of these overlapped partially or completely with a predicted gene based on the Trinity mapping or Stringtie genes. However, 3,866 genes (4,321 transcripts) had no overlap with any predicted exon from either the Trinity or StringTie set, and were kept for the final set. Considering the possibility that some of these may be pseudogenes, we aligned these proteins to the SwissProt database with BLASTP [Camacho et al., 2009]. Of these, only 759 had reliable hits (E-value < 10^−5^) to 688 proteins. The annotated functions were diverse, including proteins similar to many receptors and large structural proteins such as fibrillin (potentially any protein with EGF repeats), dynein heavy chain, and titin; because very large proteins may be split across multiple contigs, the predicted genes may be only fragments of the full gene. Only 42 of the hits were against transposable elements.

#### Filtering of the final gene set

Because assembly of transcripts for both StringTie and Trinity relies on overlaps in the genome or RNAseq reads, genes that overlap in the untranslated regions (UTRs) can sometimes be erroneously fused. For StringTie, we developed a custom Python script to separate non-overlapping transcripts belonging to the same “gene” (stringtiesplitgenes.py, available at https://bitbucket.org/wrf/sequences/). Tandem duplications can lead to RNAseq reads bridging the two tandem copies and result in both copies being called the same gene. The original StringTie set contained 46,572 transcripts for 32,112 genes, while the corrected set contained 33,200 genes and identified 1,088 new non overlapping genes.

Positional errors in the genome or allelic variations may result in some RNAseq reads not mapping to the genome, so some genes are fragmented in the genome-guided transcriptome but not the *de novo* assembly. Making use of the protein predictions from TransDecoder, we compared the predicted proteins between the two transcriptomes using a custom Python script (transdecodersplitgenes.py, available at https://bitbucket.org/wrf/sequences/). This identified 406 StringTie transcripts that were better modeled by Trinity transcripts.

#### Functional gene annotation

Many genes of functional importance were examined manually, and the best transcript from StringTie, Trinity, or AUGUSTUS was retained for the final gene set. In the GFF and fasta versions of the transcriptomes, names of protein functions were assigned several ways. Target genes that were manually curated and edited, such as those used in all trees, are named by the generic function or the annotated function of the closest human protein. For instance, the dopamine beta-hydroxylase (DBH) homolog in *T. wilhelma* was manually corrected, and the position in a phylogenetic tree demonstrated that demosponges diverged before the duplication which created DBH and the two DBH-like proteins in humans, thus the *T. wilhelma* protein is annotated as DBH-like. Secondly, automated ortholog finding pipelines (HaMStR [Ebersberger et al., 2009]) used for phylogeny [Cannon et al., 2016] have identified homologs in *T. wilhelma*, which have been manually checked based on positions in the phylogenetic trees. Thirdly, single-direction BLAST results were kept as annotations provided that the BLAST hit had a bitscore over 1000, or a bitscore over 300 and the *T. wilhelma* protein covered at least 75% of the best hit against the human protein dataset from SwissProt. The bitscore and length cutoffs were applied to reduce the number of annotations based on a single domain.

### Analysis of splice variation

Using the transcriptome from StringTie, splice variation was assessed using a custom Python script (splice-variantstats.py, available at https://bitbucket.org/wrf/sequences/). In this script, several ambiguous definitions were clarified to define the different splice types. Firstly, single exon genes with no variants are distinguished from single exon genes with variations, that is, a gene with two exons can have a variant with one exon. For loci with only two transcripts, the canonical or main transcript is defined as the one with the higher expression level, as measured by the higher FPKM value reported from StringTie. For loci with three or more transcripts, main or canonical exons are those included in at least two transcripts. A cassette exon must occur in less than 50% of the transcripts for a locus, otherwise such case is defined as a skipped exon. A retained intron is any portion that exactly spans two other exons; for highly expressed transcripts this may include erroneously retained introns due to intermediates in splicing. A summary of the splicing types is displayed in Supplemental Table 2.

Intron retention was recently reported to be a common mode of alternative splicing in *A. queens-landica* [Fernandez-Valverde et al., 2015]. We found 3,295 transcripts with 3,565 retained intron events (Supplemental Table 2). We then analyzed the length of the retained introns and found the phase of the retained piece to be randomly distributed (unlike cassette exons, Supplemental Table 3), suggesting that many of the retained introns result from incomplete splicing rather than functional retention.

### Microsynteny across sponges

Putative synteny blocks were identified using a custom Python script (microsynteny.py, available at https://bitbucket.org/wrf/sequences/). Briefly, the script combines the gene positions on scaffolds for both the query and the reference with BLASTX hits for the query against the reference. If a minimum of three genes in a row on a query scaffold match to different genes on the same reference scaffold, the group is kept. By default, this mandated a gap of no more than five genes before discarding the block, and that the next gene must occur within 30kb. This method was designed to work for highly fragmented genomes with thousands of scaffolds, so the order and direction of the corresponding genes on the reference scaffold do not need to match those of the query scaffold.

StringTie transcripts for *T. wilhelma* were aligned against the *A. queenslandica* v2.0 protein set with BLASTX [Camacho et al., 2009], and positions were taken from the accompanying *A. queenslandica* v2.0 GTF. The same procedure was attempted against the *S. ciliatum* gene models, though essentially no syntenic blocks were detected, indicating either substantial differences in gene content or gene order between demosponges and calcareous sponges.

### Collection of reference data

Proteins for *Oikopleura dioica* [Denoeud et al., 2010] were downloaded from Genoscope. Gene models for *Ciona intestinalis* [Dehal et al., 2002], *Branchiostoma floridae* [Putnam et al., 2008], *Trichoplax adherens* [Srivastava et al., 2008], *Capitella teleta, Lottia gigantea, Helobdella robusta* [Simakov et al., 2013], *Saccoglossus kowalevskii* [Simakov et al., 2015], and *Monosiga brevicollis* [King et al., 2008] were downloaded from the JGI genome portal. Gene models for *Sphaeroforma arctica, Capsaspora owczarzaki* [Suga et al., 2013] and *Salpingoeca rosetta* [Fairclough et al., 2013] were downloaded from the Broad Institute.

We used genomic data of the cnidarians *Nematostella vectensis* [Moran et al., 2014], *Exaiptasia pallida* [Baumgarten et al., 2015], and *Hydra magnipapillata* as well as transcriptomes from 33 other cnidarians [Bhattacharya et al., 2016, Zapata et al., 2015, Pratlong et al., 2015, Brinkman et al., 2015, Ponce et al., 2016], mostly corals.

For demosponges, we used the genome of *Amphimedon queenslandica* [Srivastava et al., 2010, Fernandez-Valverde et al., 2015] and transcriptomic data from: *Mycale phyllophila* [Qiu et al., 2015], *Petrosia ficiformis* [Riesgo et al., 2014a], *Crambe crambe* [Versluis et al., 2015], *Cliona varians* [Riesgo et al., 2014b], *Halisarca dujardini* [Borisenko et al., 2016], *Crella elegans* [Pérez-Porro et al., 2013], *Stylissa carteri, Xestospongia testutinaria* [Ryu et al., 2016], *Scopalina* sp., and *Tedania anhelens*. We used data from the genome of the calcareous sponge *Sycon ciliatum* [Fortunato et al., 2014] and the transcriptome of *Leucosolenia complicata*. For hexactinellids (glass sponges), we used transcriptome data from *Aphrocallistes vastus* [Ludeman et al., 2014], *Hyalonema populiferum, Rosella fibulata*, and *Sympagella nux* [Whelan et al., 2015]. For homoscleromorphs, we used two transcriptomes from *Oscarella carmela* and *Corticium candelabrum* [Ludeman et al., 2014].

We used data from the two published draft genomes of ctenophores [Ryan et al., 2013, Moroz et al., 2014], as well as transcriptome data from 11 additional ctenophores: *Bathocyroe fosteri, Bathyctena chuni, Beroe abyssicola, Bolinopsis infundibulum, Charistephane fugiens, Dryodora glandiformis, Euplokamis dunlapae, Hormiphora californensis, Lampea lactea, Thalassocalyce inconstans, and Velamen parallelum*.

We used data from the unpublished draft genome of a novel placozoan species, designated H13.

### Gene trees

For protein trees, candidate proteins were identified by reciprocal BLAST alignment using blastp or tblastn. All BLAST searches were done using the NCBI BLAST 2.2.29+ package [Camacho et al., 2009]. Because most functions were described for human, mouse, or fruit fly proteins, these served as the queries for all datasets. Candidate homologs were kept for analysis if they reciprocally aligned by blastp to a query protein, usually human. Alignments for protein sequences were created using MAFFT v7.029b, with L-INS-i parameters for accurate alignments [Katoh and Standley, 2013]. Phylogenetic trees were generated using either FastTree [Price et al., 2010] with default parameters or RAxML-HPC-PTHREADS v8.1.3 [Stamatakis, 2014], using the PROTGAMMALG model for proteins and 100 bootstrap replicates with the “rapid bootstrap” (-f a) algorithm and a random seed of 1234.

### Domain annotation

Domains for individual protein trees were annotated with “hmmscan” v3.1b1 from the HHMER package [Eddy, 2011] using the PFAM-A database v27.0 [Finn et al., 2016] as queries. Signal peptides were predicted using the stand-alone version of SignalP v4.1 [Petersen et al., 2011]. Domain structures were visualized using a custom Python script, “pfampipeline.py”, available at https://github.com/wrf/genomeGTFtools.

## Acknowledgments

W.R.F would like to thank K. Achim, M. Nickel, J. Musser, J. Ryan, and I. Oldenburg for helpful comments. G.W and D.E. would like to thank M. Nickel for providing the initial *T. wilhelma* specimens to set up the culture in Munich. This work was supported by a LMUexcellent grant (Project MODELSPONGE) to D.E. and G.W. as part of the German Excellence Initiative, partially by research grant 9278 (“Early evolution of multicellular sponges”) from VILLUM FONDEN to G.W., and NIH grant NIGMS-5-R01-GM087198 to S.H.D.H. The authors declare no competing interests.

## 1 Supplemental Figures

**Supplemental Figure 1:**
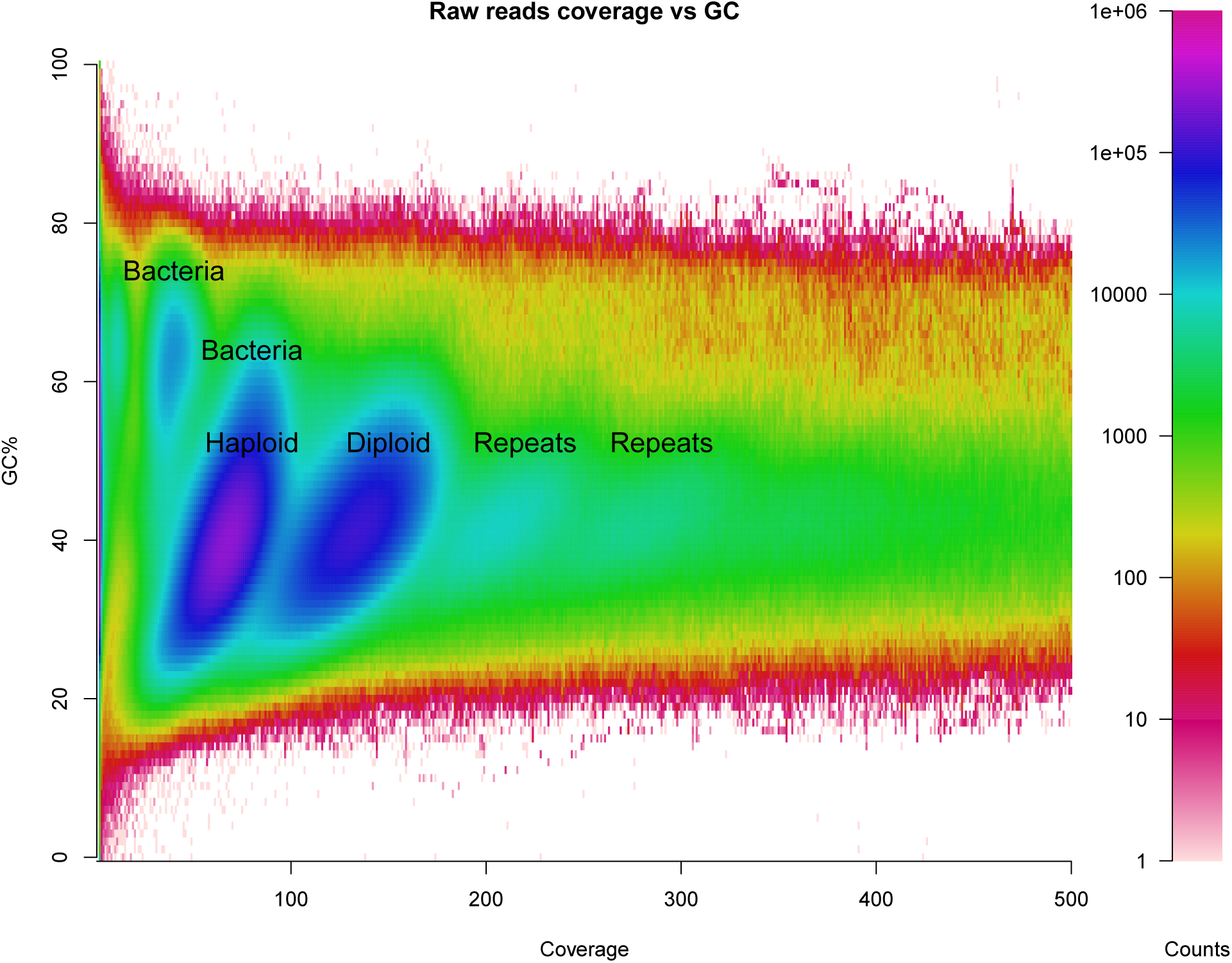
Coverage vs. GC content for reads. Lava lamp plot of the unfiltered paired-end reads. Coverage was calculated as the median 31-mer coverage for each read. High-GC reads indicate the presence of bacteria in the raw reads.

**Supplemental Figure 2:**
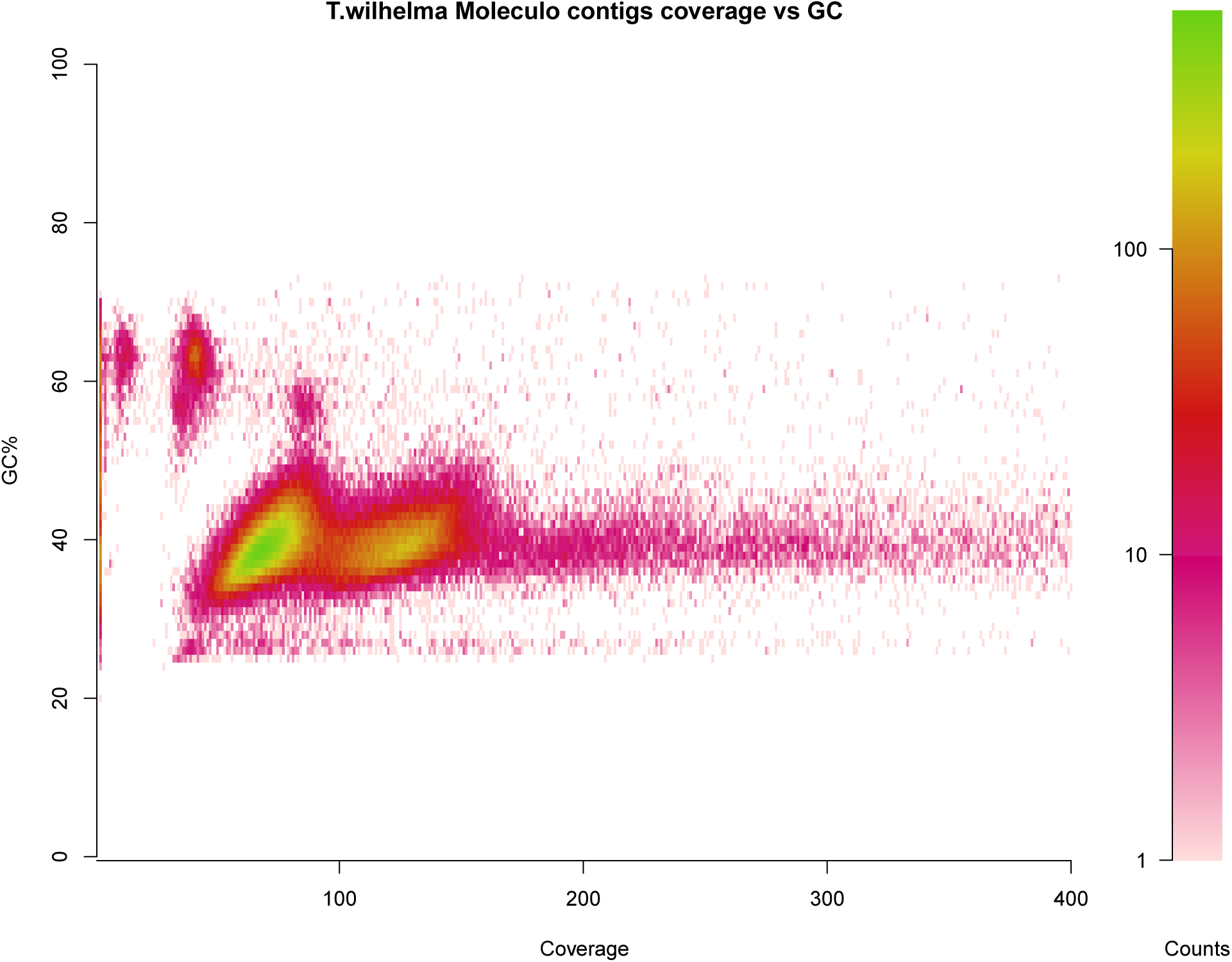
Coverage vs. GC content for genomic scaffolds. Heat map of percent GC versus coverage of reads for the all Moleculo reads.

**Supplemental Figure 3:**
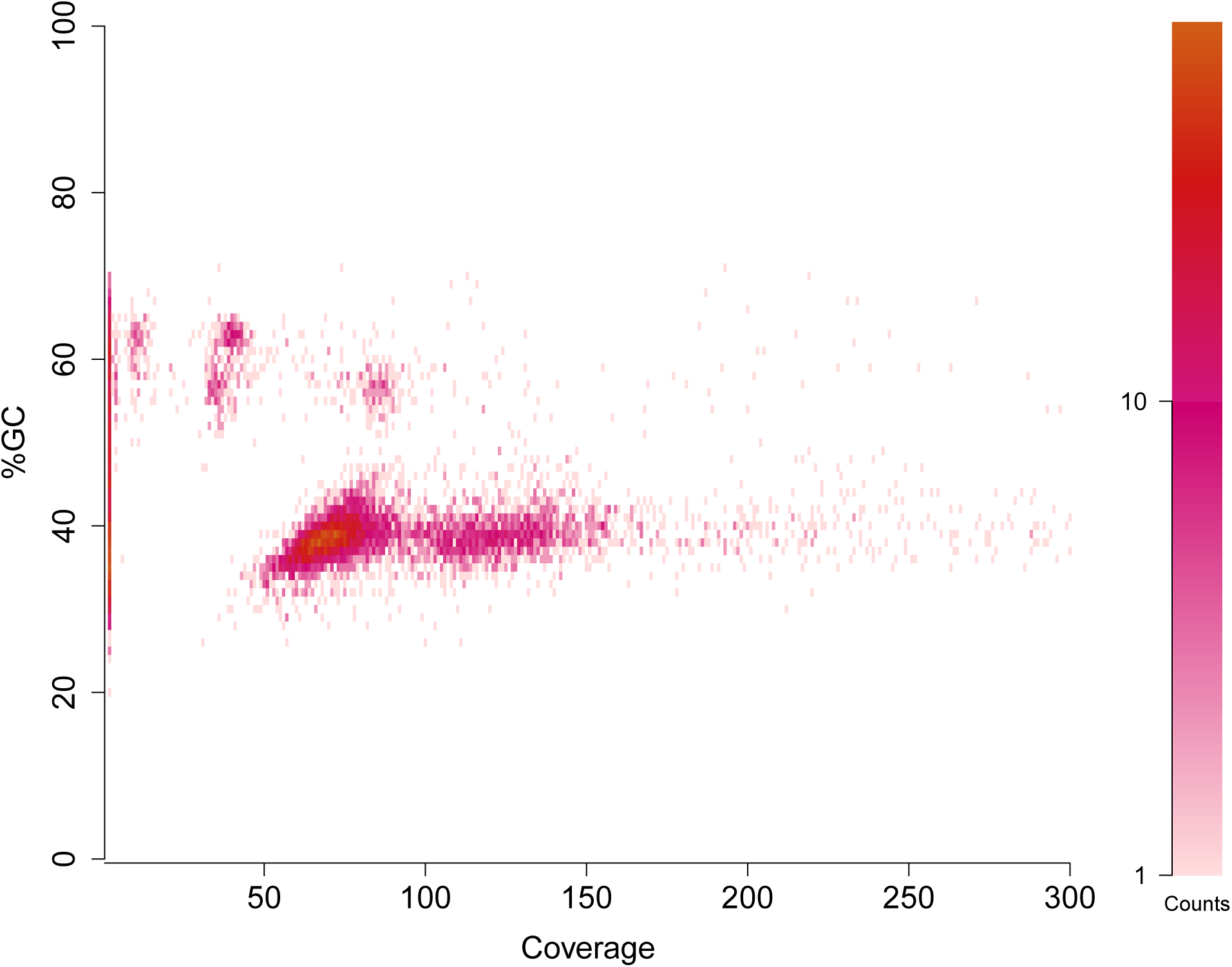
Coverage vs. GC content for genomic scaffolds. Scatterplot of percent GC versus coverage of reads for the all scaffolds. The 1,040 contigs with zero coverage are carried over from low-coverage Moleculo reads, and likely derive from amplified contaminating DNA.

**Supplemental Figure 4:**
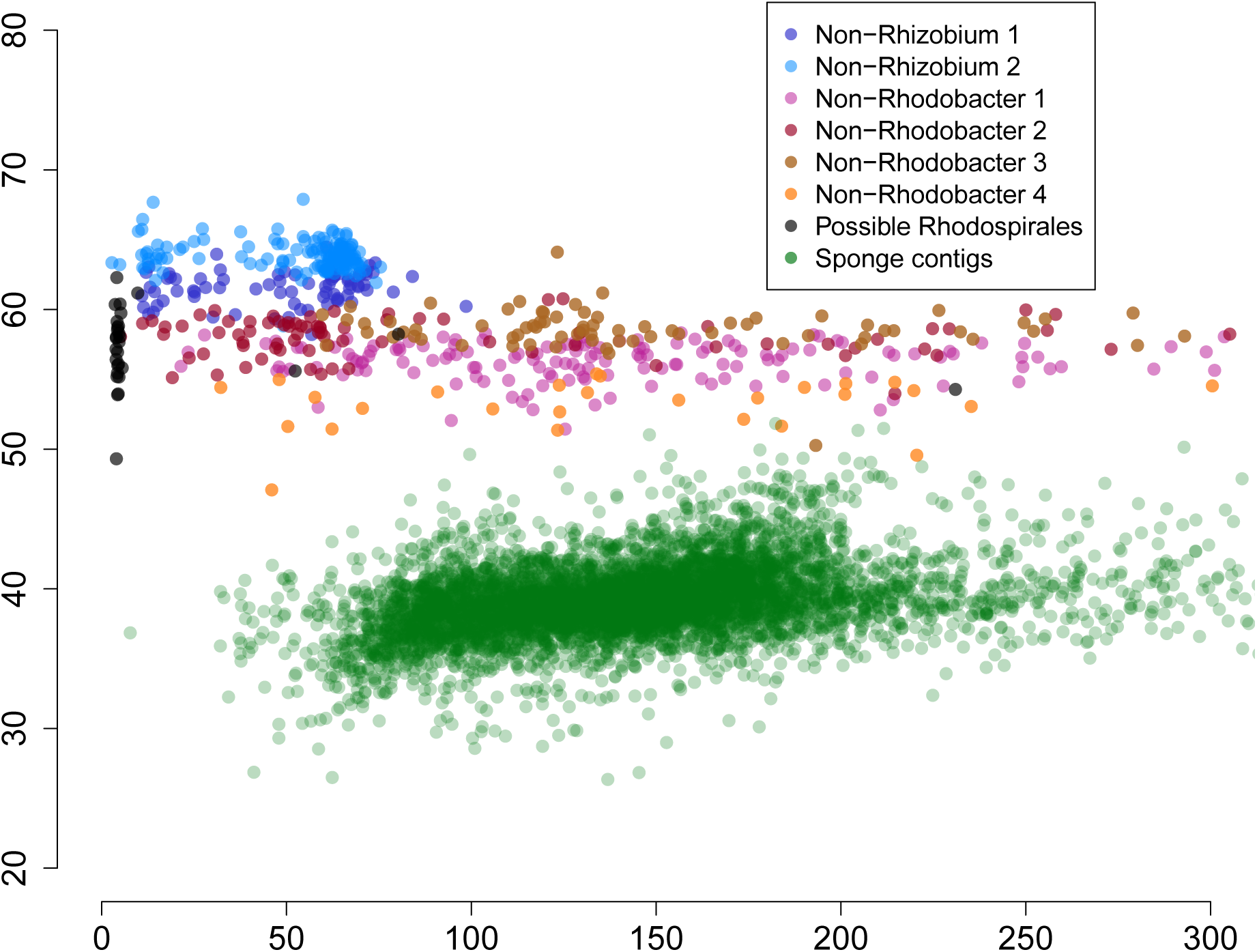
Separation of contigs derived from bacterial symbionts. Sponge scaffolds are in green, while scaffolds assigned to bacteria are identified by blue (*Rhizobiales*) and pink (*Rhodobacter*). Low-coverage contigs were removed. Seven bins were identified with MetaWatt to separate the bacterial contigs, though several bins appeared to correspond to the same bacteria.

**Supplemental Figure 5:**
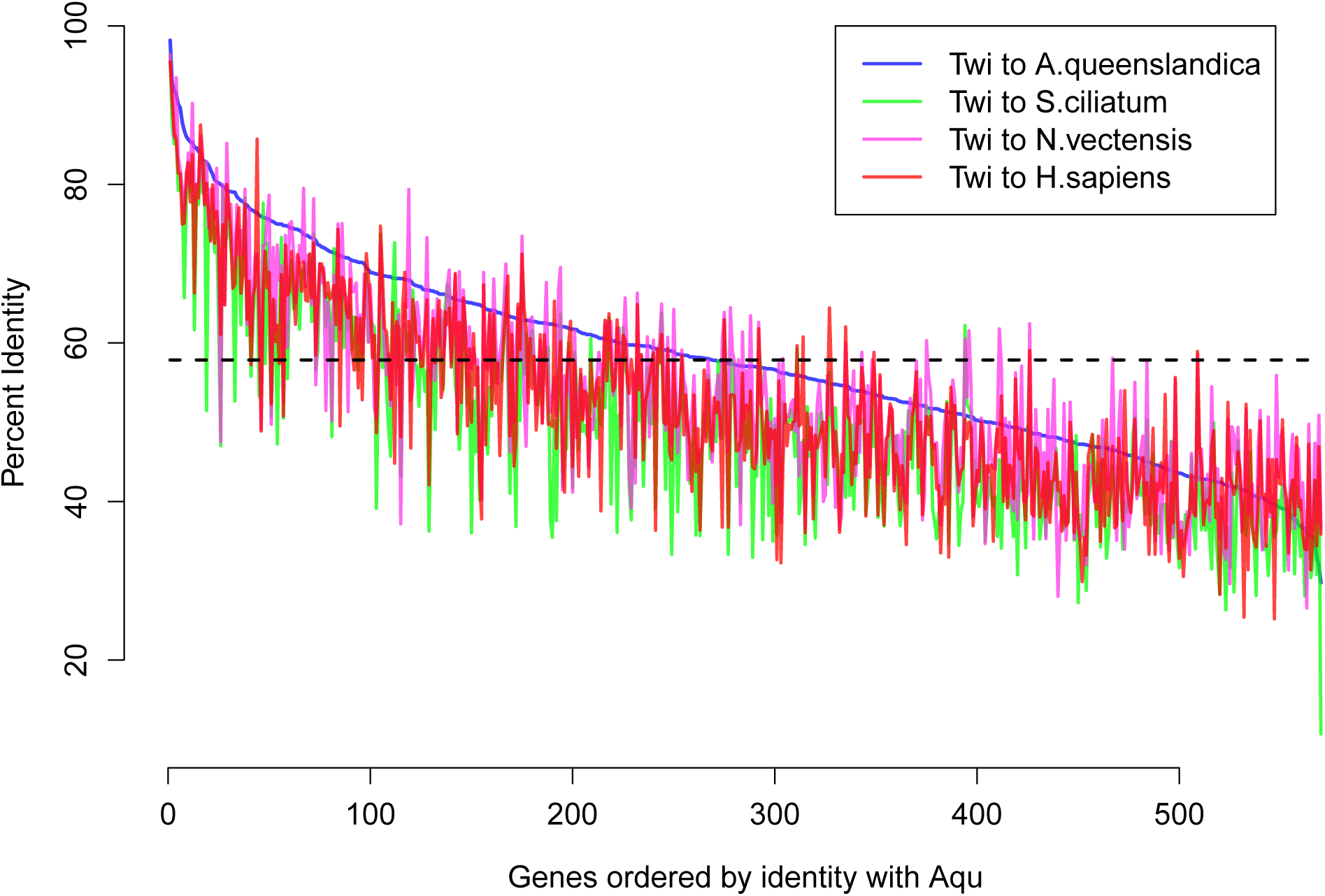
Percent identity between sponges. Calculated protein percent identity between 570 one-to-one orthologs between *T. wilhelma* and *A. queens-landica* (blue), *S. ciliatum* (green), the anemone *N. vectensis* (purple) and human (red). Average identity between *T. wilhelma* and *A. queenslandica* is shown as the dotted black line at 57%.

**Supplemental Figure 6:**
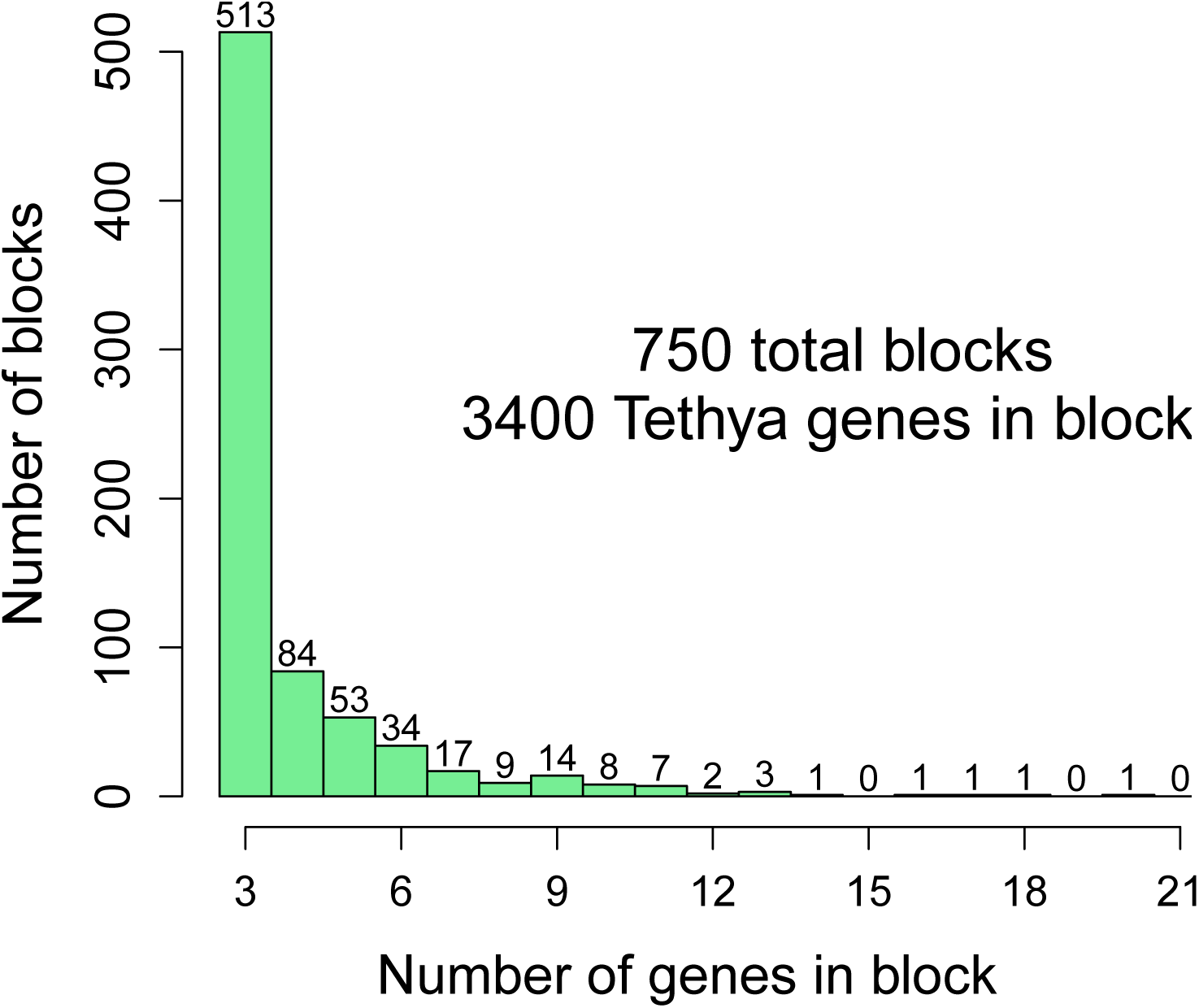
Length of microsyntenic blocks. Histogram of number of genes in detected microsyntenic blocks between *T. wilhelma* and *A. queenslandica* v2.0 gene models.

**Supplemental Figure 7:**
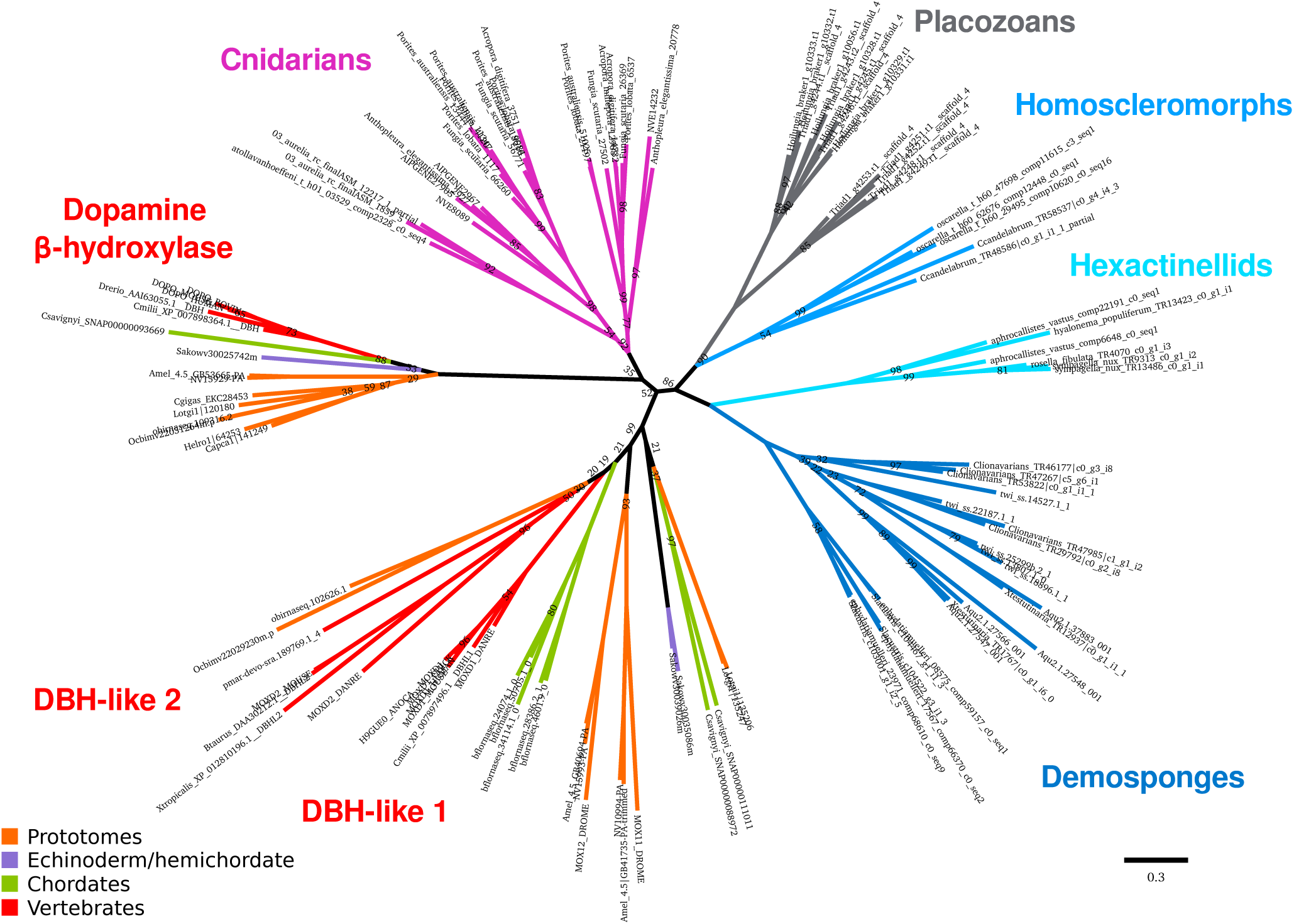
Dopamine *β*-hydroxylase homologs across metazoans. Tree of DBH and DBH-like proteins across all metazoan groups, generated with RAxML using the PROTGAMMALG model. Bootstrap values are 100 unless otherwise shown.

**Supplemental Figure 8:**
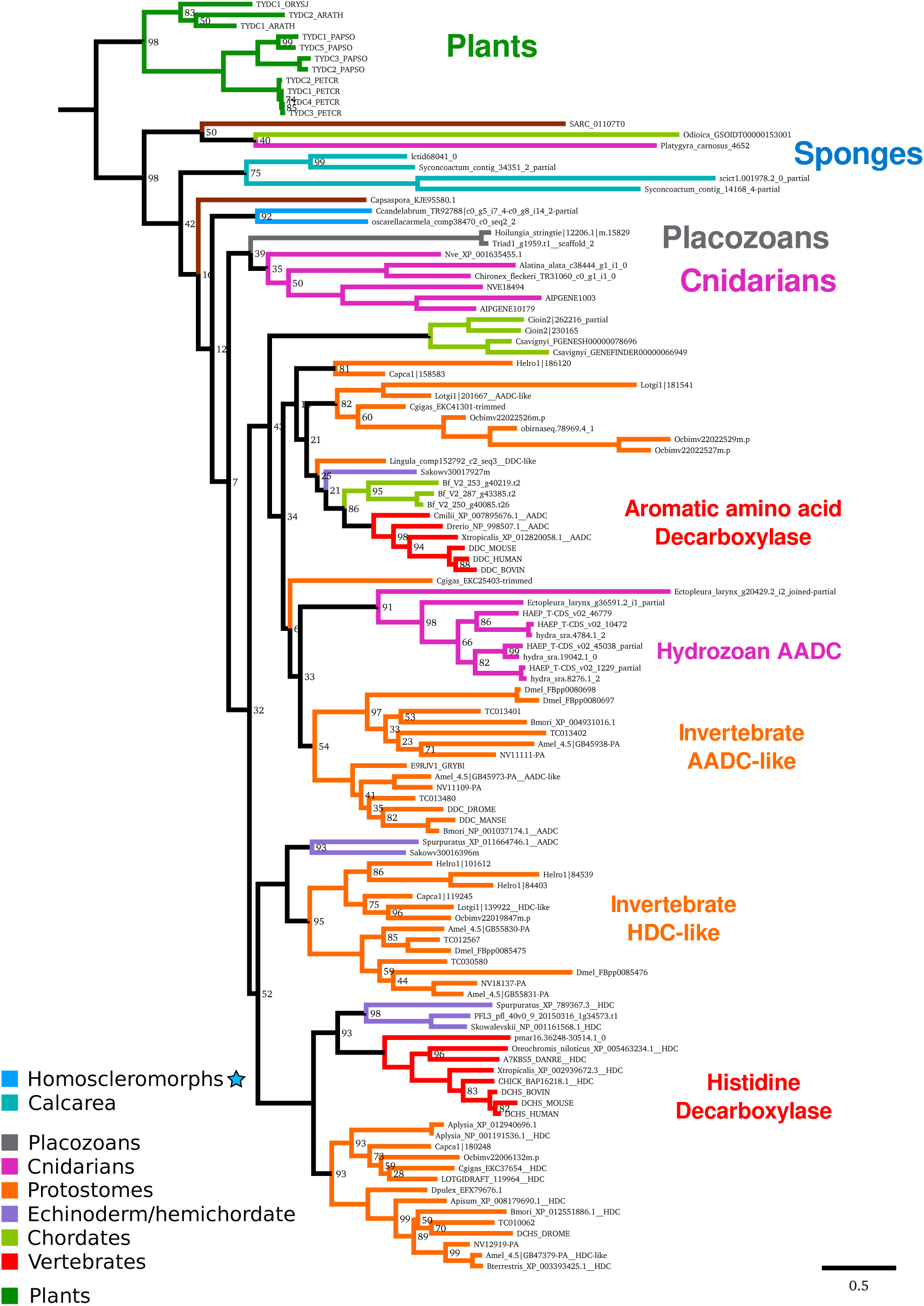
Aromatic amino acid decarboxylase homologs across metazoans. Tree of AADC and histidine decarboxylase (HDC) proteins across all metazoan groups, generated with RAxML using the PROTGAMMALG model. Bootstrap values are 100 unless otherwise shown.

**Supplemental Figure 9:**
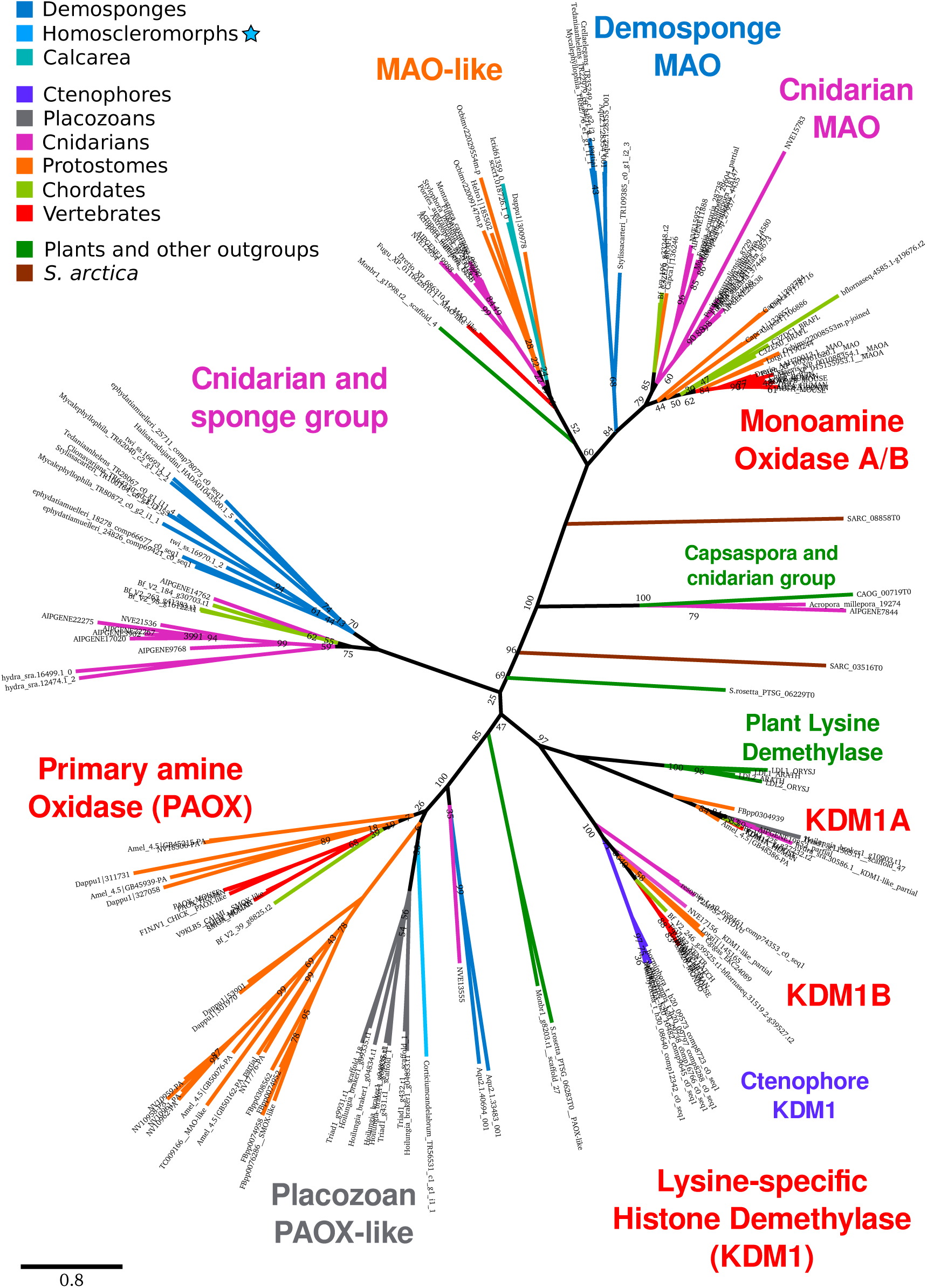
Monoamine oxidase homologs across metazoans. Tree of monoamine oxidase (MAO) and related proteins across all metazoan groups, generated with RAxML using the PROTGAMMALG model. Searches for lysine demethylase (KDM1) and primary amine oxidase (PAOX) were not exhaustive, and were added to display the sole positions of ctenophores (only have KDM) and placozoans (only have PAOX) in this protein family. Bootstrap values are 100 unless otherwise shown.

**Supplemental Figure 10:**
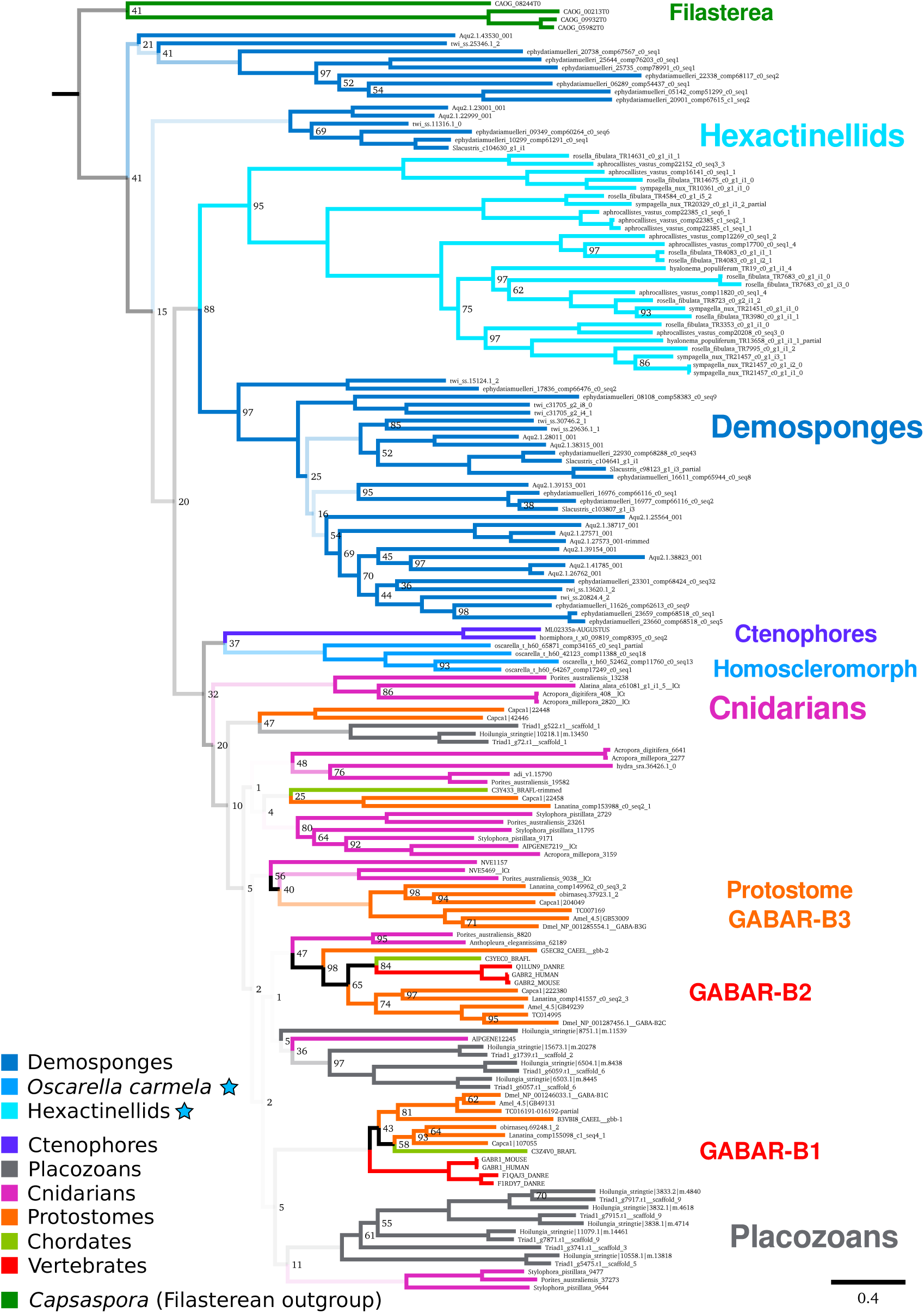
mGABA receptors across metazoans. Complete version of Figure 4 metabotropic GABA receptor (GABA-B type) protein tree generated with RAxML. Bootstrap values are 100 unless otherwise shown. The majority of deeper nodes were poorly resolved; branch transparency corresponds to bootstrap support for values under 50, meaning half-transparent.

**Supplemental Figure 11:**
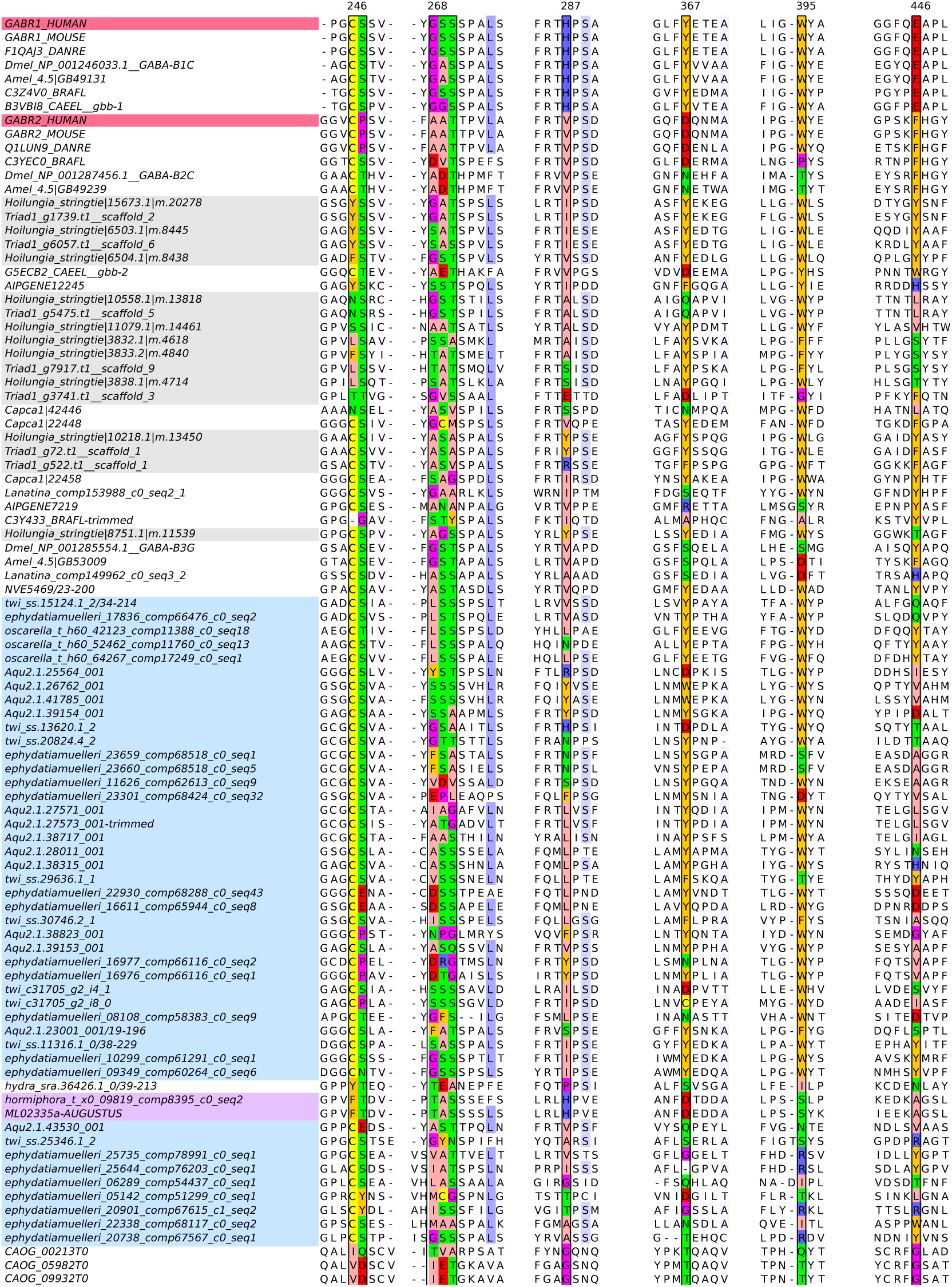
Binding pocket alignment of mGABARs across metazoans. Select residues involved in the binding of GABA are highlighted, and numbers correspond to the position in the human protein GABR1/GABAR-B1, based on the structure of GABR1 [Geng et al., 2013]. Highly conserved residues not thought to be involved in binding are highlighted in blue. The two human proteins are highlighted in pink. Placozoan proteins are highlighted in gray. Sponge proteins are highlighted in blue. The two ctenophore proteins are highlighted in violet.

**Supplemental Figure 12:**
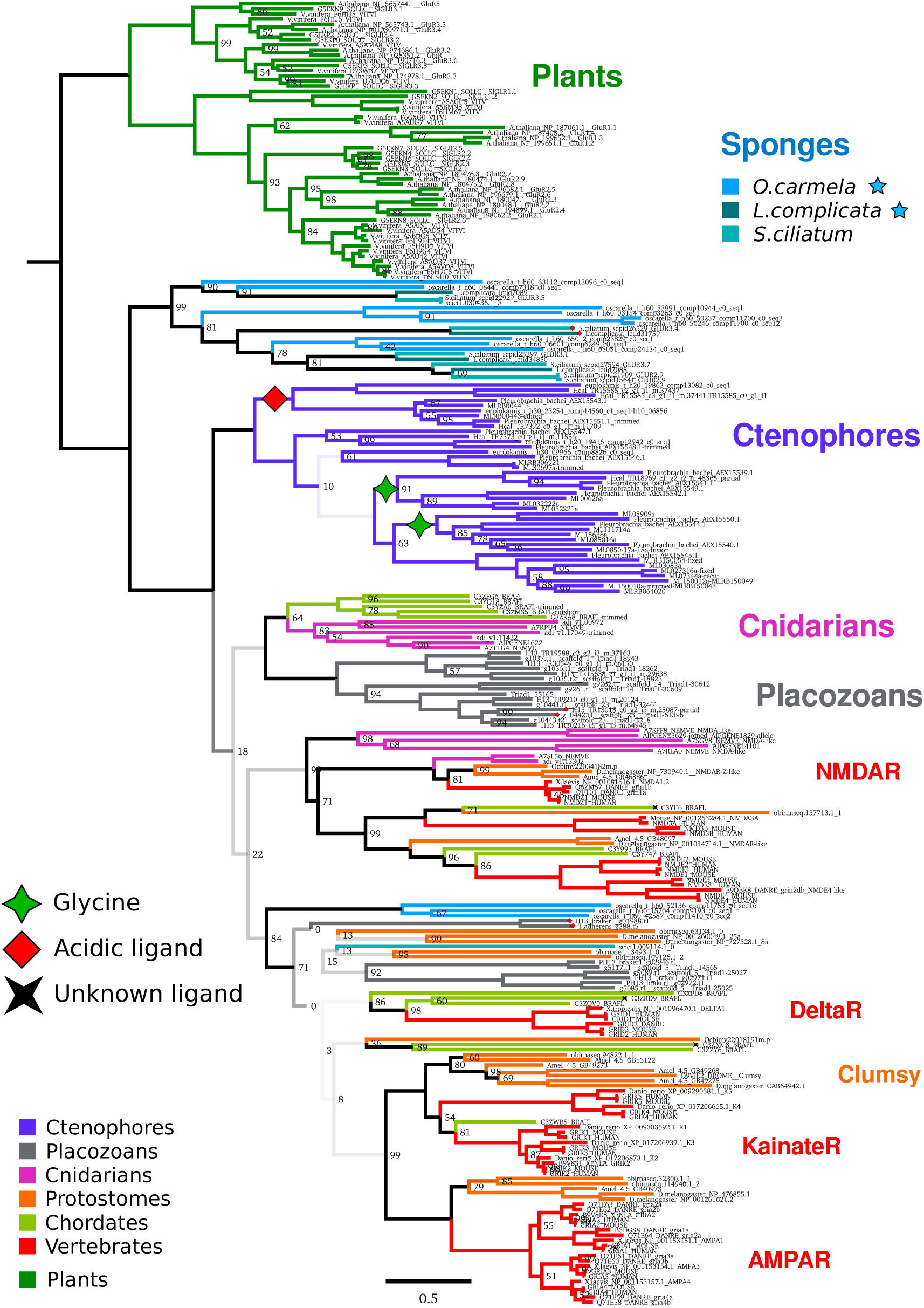
Phylogenetic tree of ionotropic glutamate receptors across metazoans. Protein tree generated with RAxML using the PROTCATWAG model. Bootstrap values are 100 unless otherwise shown. These receptors are not found in the genomes or transcriptomes of demosponges or hexactinellids, so the Sponge clade refers to calcareous sponges and homoscleromorphs. For the three sponges, the blue star indicates sequences derived from a transcriptome. Based on [Alberstein et al., 2015], some receptors are predicted to bind ligands other than glutamate, shown with the green star, red diamond, and black star, for glycine, acidic ligands, and unknown, respectively. Four placozoan proteins have substitutions at the conserved acidic residue (D723 in human GluN1), as either GY in ctenophores, or GG/WY in placozoans; the carboxyl of the glutamic/aspartic acid is needed to coordinate the amino group of glutamate, suggesting that these proteins do not bind an *α*-amino acid.

**Supplemental Figure 13:**
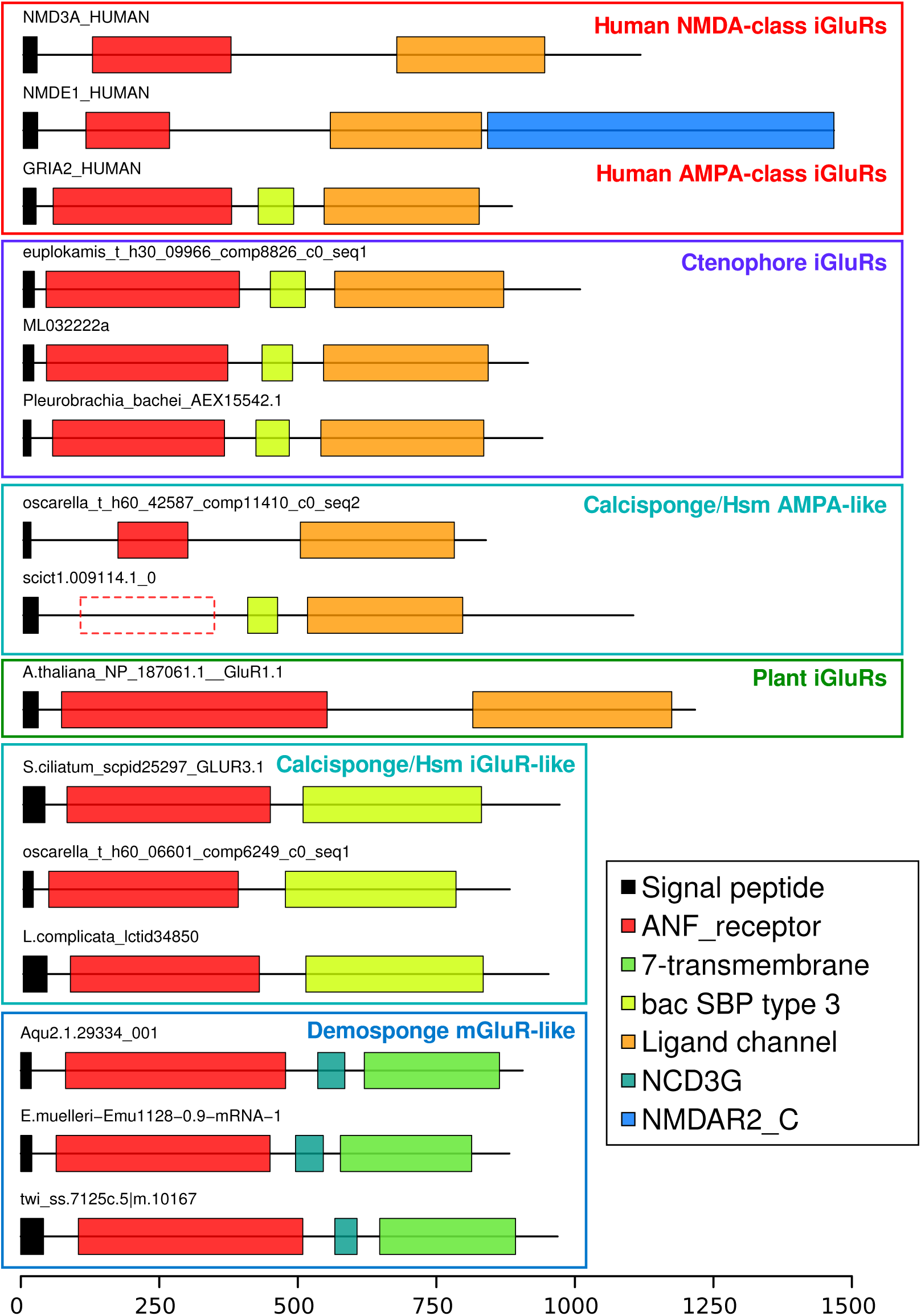
Domain organization of ionotropic glutamate receptors across metazoans. Scale bar displays number of amino acids. Top BLAST hits for human iGluRs in demosponges appear to be metabotropic, due to the presence of a 7-transmembrane domain instead of the ion channel, while the ligand-binding domain is conserved. Ctenophore iGluRs and some calcarea/homoscleromorph (Hsm) proteins have the vertebrate-type domain organization, though the other calcarea/homoscleromorph proteins (main sponge group in Supplemental Figure 12) have an SBP domain.

**Supplemental Figure 14:**
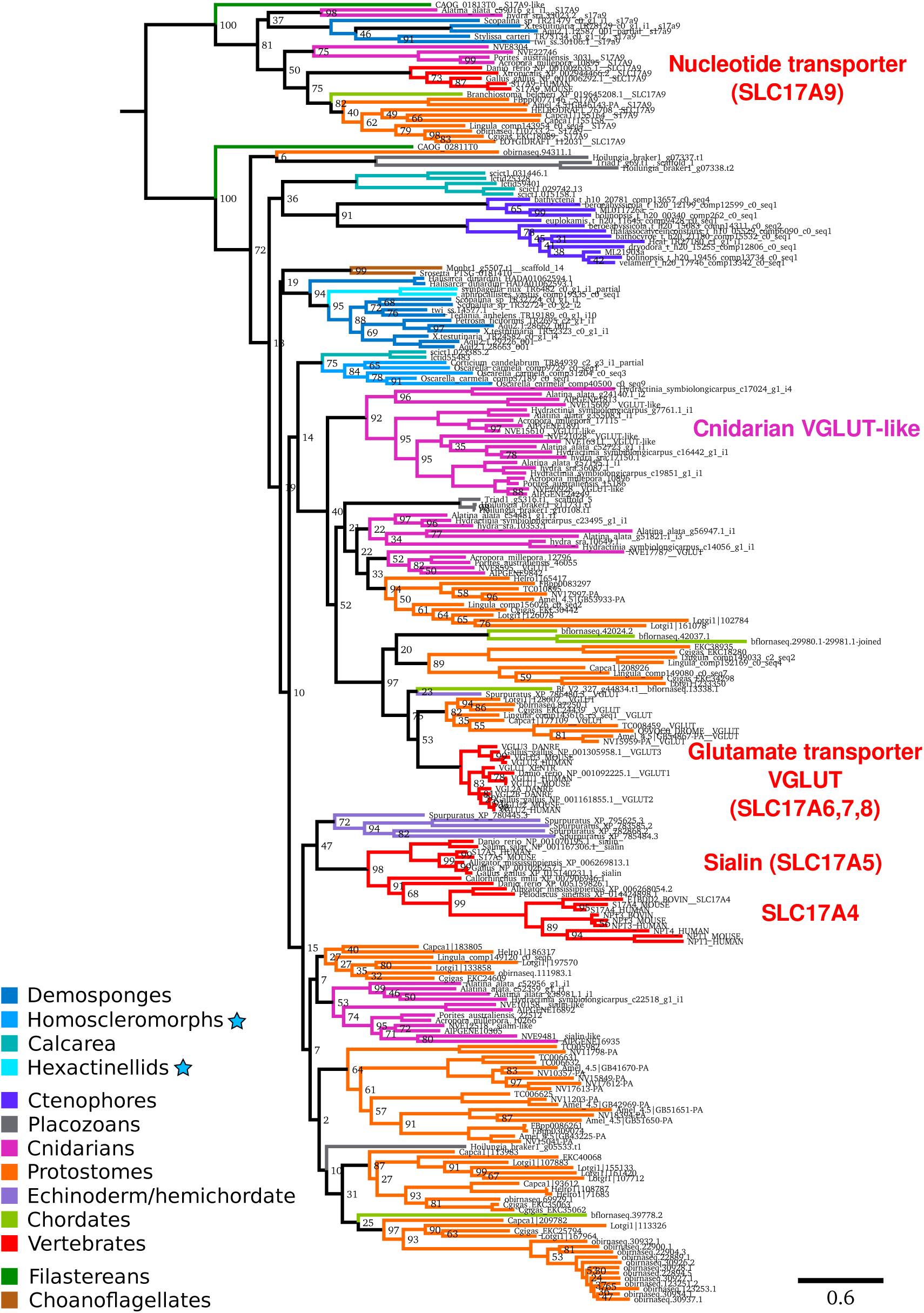
Vesicular glutamate transporter homologs across metazoans. Tree of VGLUT (SLC17A6-8) proteins across all metazoan groups, generated with RAxML using the PROTGAMMALG model. Bootstrap values are 100 unless otherwise shown.

**Supplemental Figure 15:**
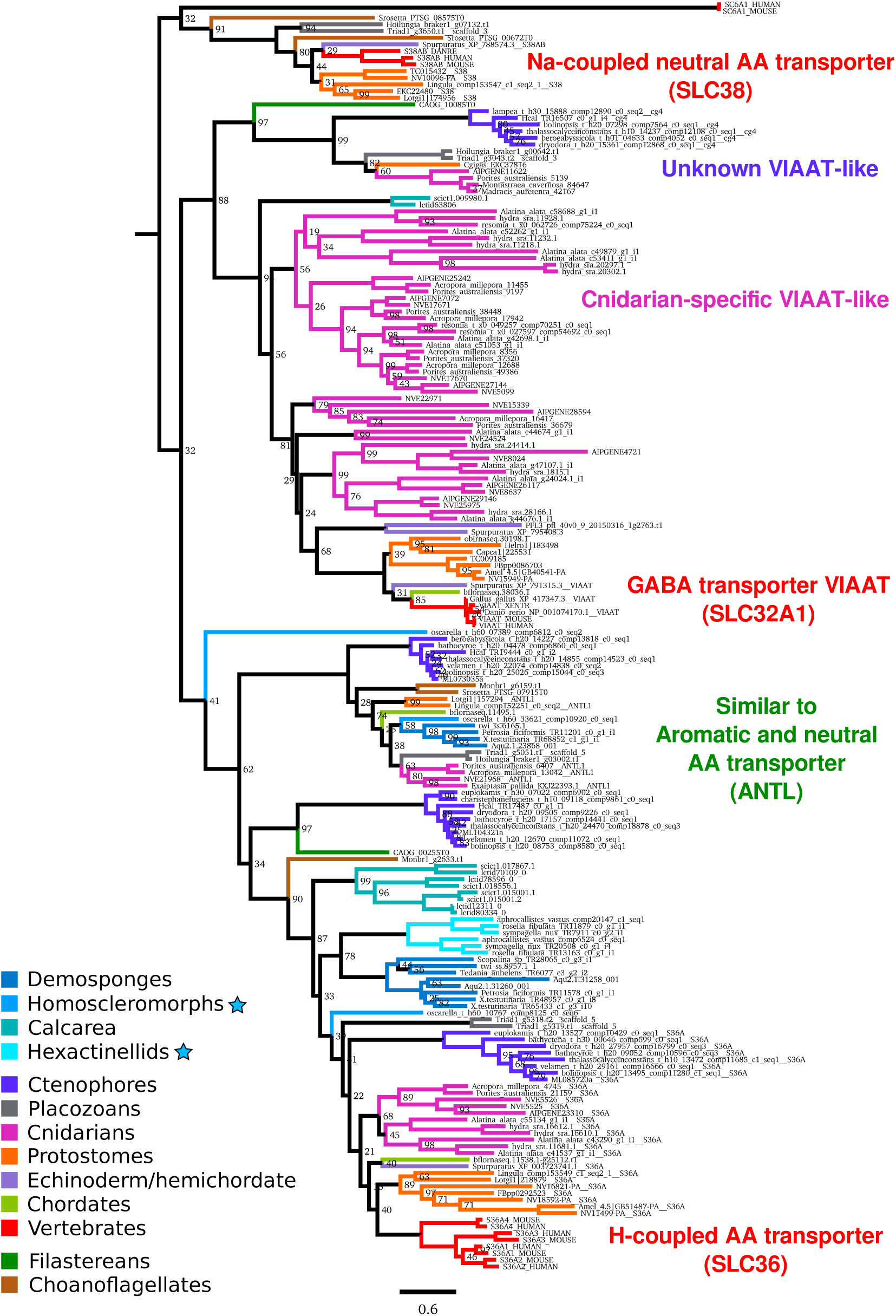
Vesicular inhibitory amino acid transporter homologs across metazoans. Tree of VIAAT (SLC32A1) proteins and related transpoters across all metazoan groups, generated with RAxML using the PROTGAMMALG model. Bootstrap values are 100 unless otherwise shown.

**Supplemental Figure 16:**
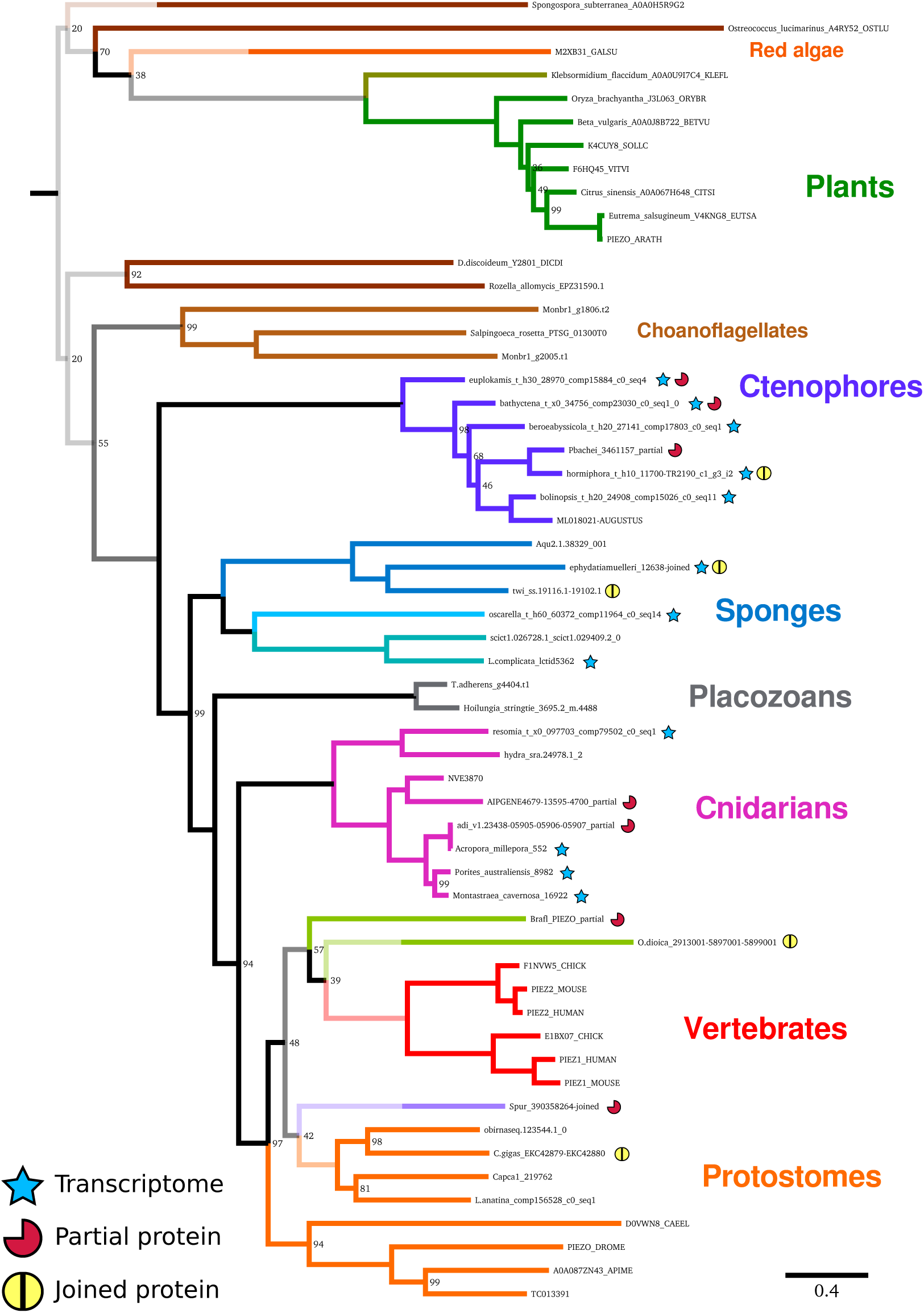
Phylogenetic tree of Piezo homologs across metazoans. Protein tree generated with RAxML using the PROTCATWAG model and 100 bootstraps. Yellow circle indicates the sequence was compete when joined with other genes or partial sequences, red partial circle indicates the sequence is incomplete in the genome or transcriptome. A blue star indicates that the sequence derived from a transcriptome, so copy number cannot be determined with certainty. Bootstrap values are 100 unless otherwise shown.

**Supplemental Figure 17:**
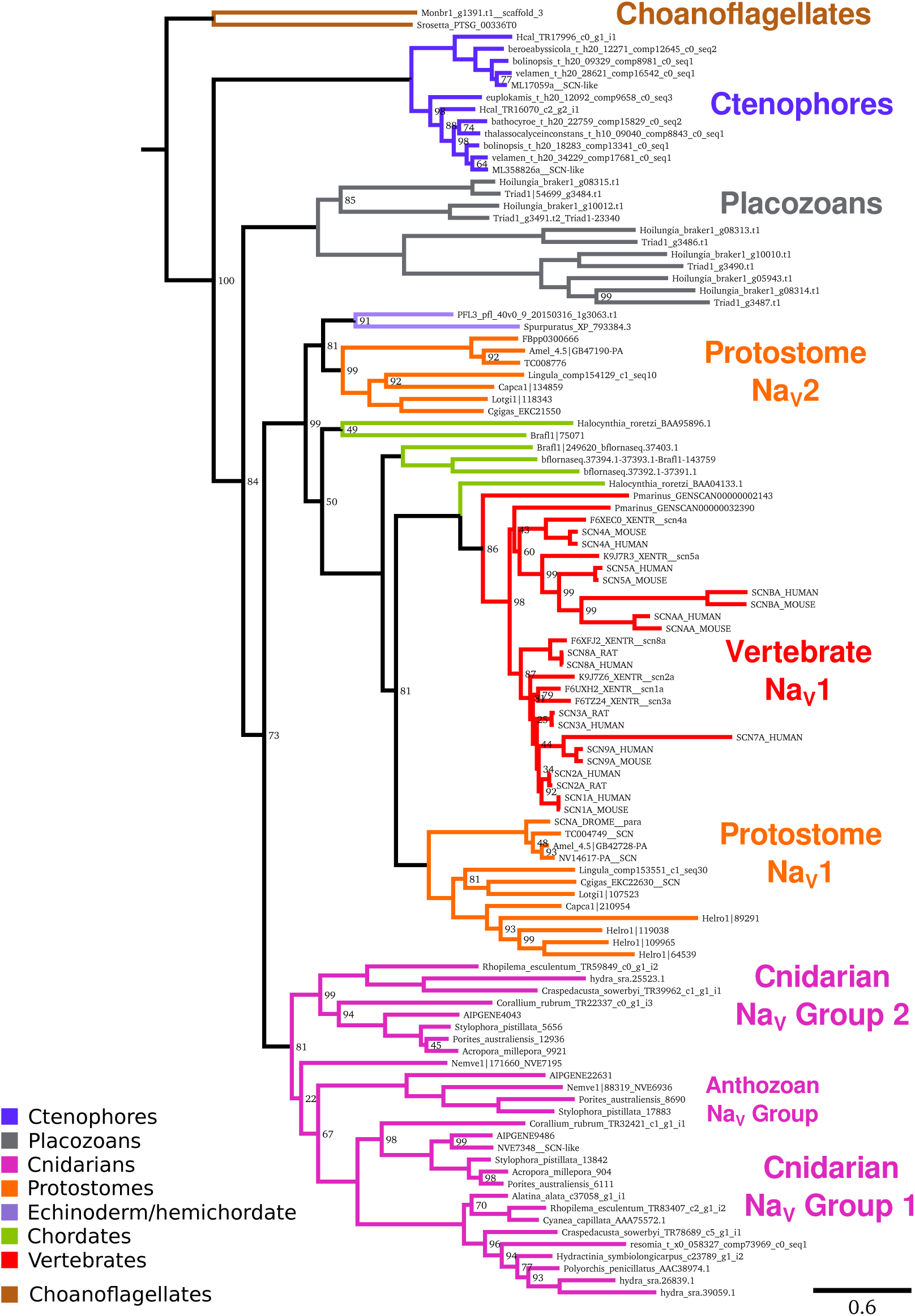
Phylogenetic tree of voltage-gated sodium channel alpha subunits across metazoans. Protein tree generated with RAxML using the PROTGAMMALG model and 100 bootstraps. Bootstrap values are 100 unless otherwise shown.

**Supplemental Figure 18:**
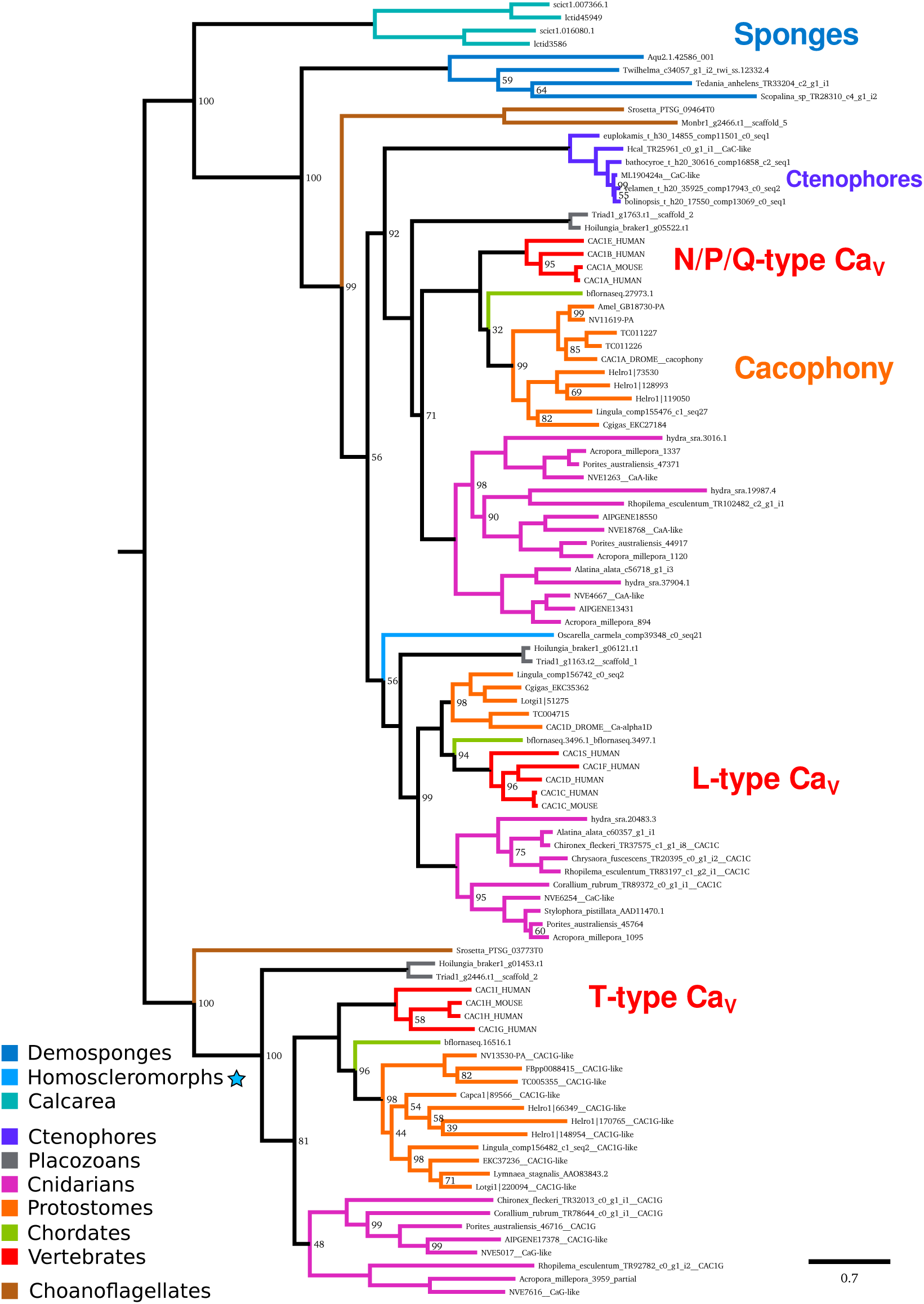
Phylogenetic tree of voltage-gated calcium channels across metazoans. Protein tree generated with RAxML using the PROTGAMMALG model and 100 bootstraps. Bootstrap values are 100 unless otherwise shown.

**Supplemental Figure 19:**
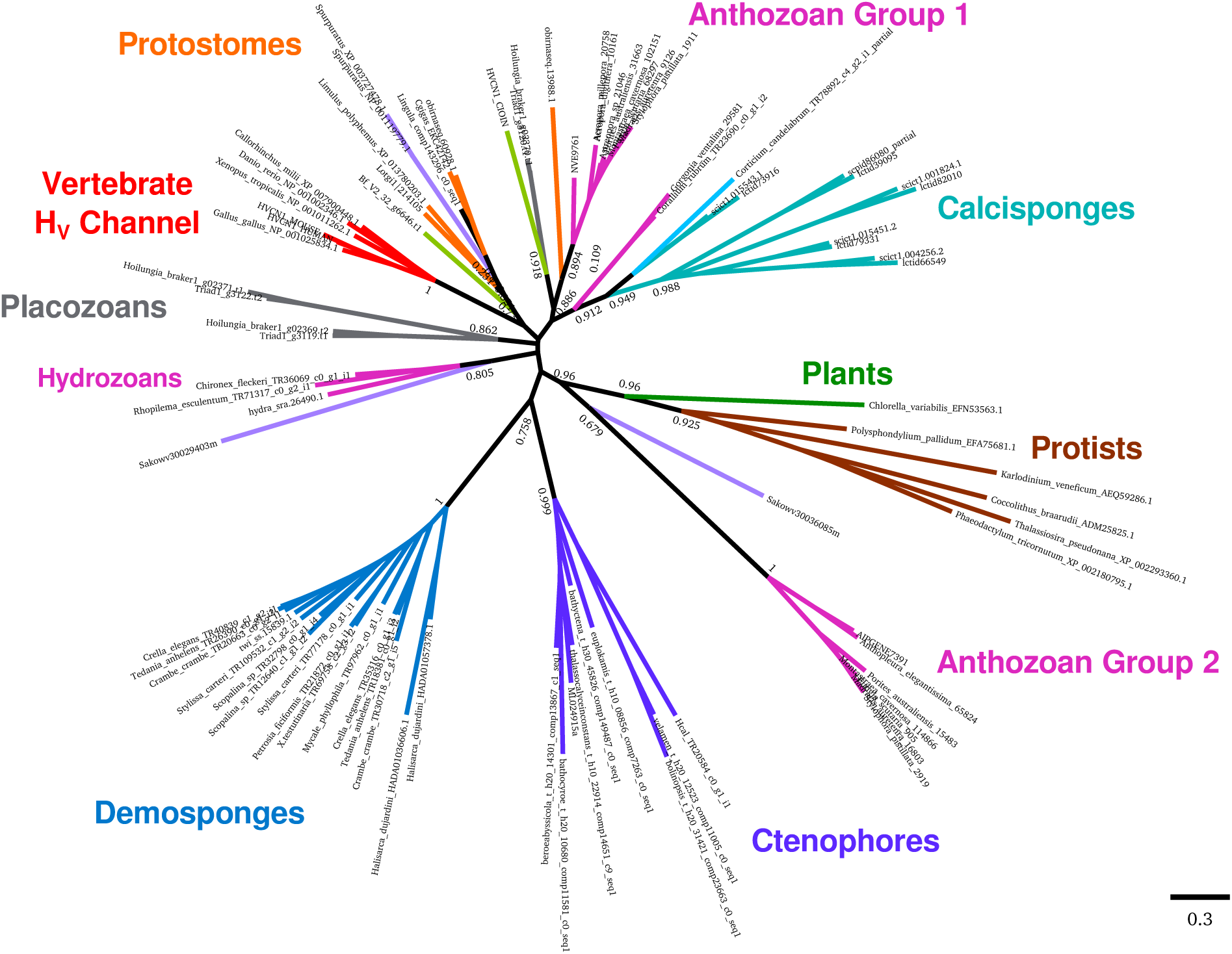
Phylogenetic tree of voltage-gated proton channels across metazoans. Protein tree generated with FastTree.

## 2 Supplemental Tables

**Supplemental Table 1:**
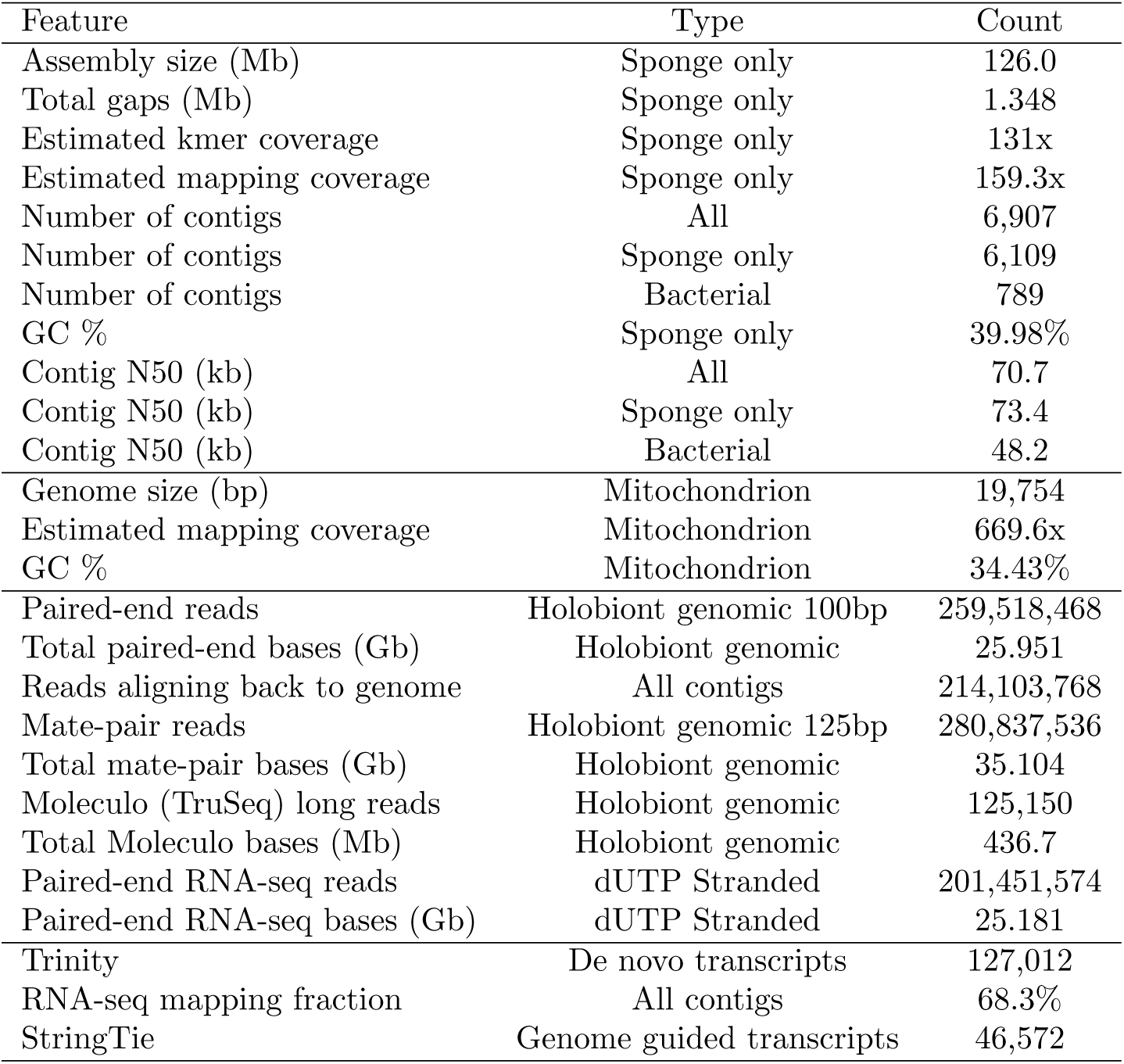
Summary statistics of the *Tethya wilhelma* genome. Holobiont genomic includes the sponge and all associated bacteria.

**Supplemental Table 2:** Summary of splice variation for the genome-guided transcriptome for *T. wilhelma* (Twi) and transcript set v2.0 for *A. queenslandica* (Aqu).

| Splice Type | Twi transcripts | Twi events/exons | Aqu transcripts | Aqu events/exons |
| --- | --- | --- | --- | --- |
| Cassette exons | 4779 | 9089 | 3591 | 5535 |
| Canonical splicing | 5049 | - | 2602 | - |
| Skipped exons | 3868 | 8329 | 1022 | 721 |
| Alternative splice acceptor | - | 3747 | - | 638 |
| Intron retention | 3295 | 3565 | 3437 | 3400 |
| Alternative splice donor | - | 3264 | - | 521 |
| Alternative N-terminus | 1964 | - | 1965 | - |
| Alternative C-terminus | 1788 | - | 1968 | - |
| Intronic start | 471 | - | 571 | - |
| Intronic end | 285 | - | 592 | - |
| Non-canonical | 73 | - | 135 | - |
| Single exon with variants | 246 | - | 13 | - |
| No variants | 12088 | - | 24027 | - |
| Single exon and no variants | 15421 | - | 12400 | - |

**Supplemental Table 3:**
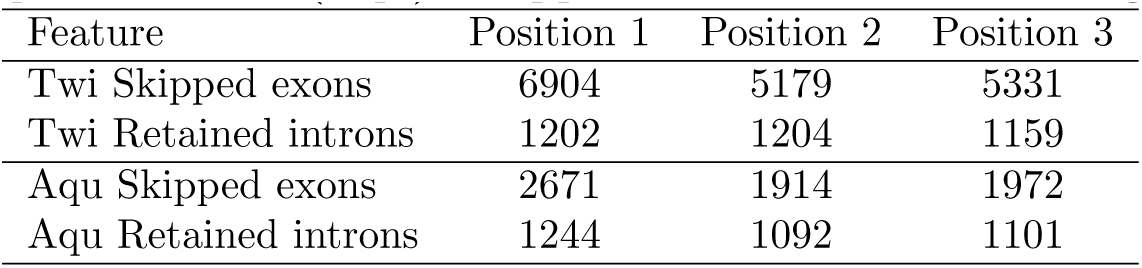
Skipped exon and retained intron frame, for *T. wilhelma* (Twi) and *A. queenslandica* (Aqu). Skipped exons tend to have lengths as multiples of three.

| Feature | Position 1 | Position 2 | Position 3 |
| --- | --- | --- | --- |
| Twi Skipped exons | 6904 | 5179 | 5331 |
| Twi Retained introns | 1202 | 1204 | 1159 |
| Aqu Skipped exons | 2671 | 1914 | 1972 |
| Aqu Retained introns | 1244 | 1092 | 1101 |

## References

Alberstein, R., Grey, R., Zimmet, A., Simmons, D. K., and Mayer, M. L. (2015). Glycine activated ion channel subunits encoded by ctenophore glutamate receptor genes. Proceedings of the National Academy of Sciences, 112(44):E6048–E6057.

Árnadóttir, J. and Chalfie, M. (2010). Eukaryotic Mechanosensitive Channels. Annual Review of Biophysics, 39(1):111–137.

Baumgarten, S., Simakov, O., Esherick, L. Y., Liew, Y. J., Lehnert, E. M., Michell, C. T., Li, Y., Hambleton, E. a., Guse, A., Oates, M. E., Gough, J., Weis, V. M., Aranda, M., Pringle, J. R., and Voolstra, C. R. (2015). The genome of Aiptasia, a sea anemone model for coral symbiosis. Proceedings of the National Academy of Sciences, page 201513318.

Bhattacharya, D., Agrawal, S., Aranda, M., Baumgarten, S., Belcaid, M., Drake, J. L., Erwin, D., Foret, S., Gates, R. D., Gruber, D. F., Kamel, B., Lesser, M. P., Levy, O., Liew, Y. J., MacManes, M., Mass, T., Medina, M., Mehr, S., Meyer, E., Price, D. C., Putnam, M., Qiu, H., Shinzato, C., Shoguchi, E., Stokes, A. J., Tambutté, S., Tchernov, D., Voolstra, C. R., Wagner, N., Walker, C. W., Weber, A. P., Weis, V., Zelzion, E., Zoccola, D., and Falkowski, P. G. (2016). Comparative genomics explains the evolutionary success of reef-forming corals. eLife, 5: 1–26.

Boetzer, M., Henkel, C. V., Jansen, H. J., Butler, D., and Pirovano, W. (2011). Scaffolding pre-assembled contigs using SSPACE. Bioinformatics, 27(4): 578–579.

Borisenko, I., Adamski, M., Ereskovsky, A., and Adamska, M. (2016). Surprisingly rich repertoire of Wnt genes in the demosponge Halisarca dujardini. BMC evolutionary biology, 16(1):123.

Brinkman, D. L., Jia, X., Potriquet, J., Kumar, D., Dash, D., Kvaskoff, D., and Mulvenna, J. (2015). Transcriptome and venom proteome of the box jellyfish Chironex fleckeri. BMC Genomics, 16(1):407.

Camacho, C., Coulouris, G., Avagyan, V., Ma, N., Papadopoulos, J., Bealer, K., and Madden, T. L. (2009). BLAST+: architecture and applications. BMC bioinformatics, 10: 421.

Cannon, J. T., Vellutini, B. C., Smith, J., Ronquist, F., Jondelius, U., and Hejnol, A. (2016). Xenacoelomorpha is the sister group to Nephrozoa. Nature, 530(7588):89–93.

Cao, J., Shi, F., Liu, X., Huang, G., and Zhou, M. (2010). Phylogenetic analysis and evolution of aromatic amino acid hydroxylase. FEBS Letters, 584(23): 4775–4782.

Carlberg, M. and Rosengren, E. (1985). Biochemical basis for adrenergic neurotransmission in coelenterates. Journal of Comparative Physiology B, 155(2): 251–255.

Coste, B., Xiao, B., Santos, J. S., Syeda, R., Grandl, J., Spencer, K. S., Kim, S. E., Schmidt, M., Mathur, J., Dubin, A. E., Montal, M., and Patapoutian, A. (2012). Piezo proteins are pore-forming subunits of mechanically activated channels. Nature, 483(7388):176–181.

Dehal, P., Satou, Y., Campbell, R. K., Chapman, J., Degnan, B., De Tomaso, A., Davidson, B., Di Gregorio, A., Gelpke, M., Goodstein, D. M., Harafuji, N., Hastings, K. E. M., Ho, I., Hotta, K., Huang, W., Kawashima, T., Lemaire, P., Martinez, D., Meinertzhagen, I. a., Necula, S., Nonaka, M., Putnam, N., Rash, S., Saiga, H., Satake, M., Terry, A., Yamada, L., Wang, H.-G., Awazu, S., Azumi, K., Boore, J., Branno, M., Chin-Bow, S., DeSantis, R., Doyle, S., Francino, P., Keys, D. N., Haga, S., Hayashi, H., Hino, K., Imai, K. S., Inaba, K., Kano, S., Kobayashi, K., Kobayashi, M., Lee, B.-I., Makabe, K. W., Manohar, C., Matassi, G., Medina, M., Mochizuki, Y., Mount, S., Morishita, T., Miura, S., Nakayama, A., Nishizaka, S., Nomoto, H., Ohta, F., Oishi, K., Rigoutsos, I., Sano, M., Sasaki, A., Sasakura, Y., Shoguchi, E., Shin-i, T., Spagnuolo, A., Stainier, D., Suzuki, M. M., Tassy, O., Takatori, N., Tokuoka, M., Yagi, K., Yoshizaki, F., Wada, S., Zhang, C., Hyatt, P. D., Larimer, F., Detter, C., Doggett, N., Glavina, T., Hawkins, T., Richardson, P., Lucas, S., Kohara, Y., Levine, M., Satoh, N., and Rokhsar, D. S. (2002). The draft genome of Ciona intestinalis: insights into chordate and vertebrate origins. Science (New York, N.Y.), 298(5601):2157–2167.

Denoeud, F., Henriet, S., Mungpakdee, S., Aury, J.-M., Da Silva, C., Brinkmann, H., Mikhaleva, J., Olsen, L. C., Jubin, C., Canestro, C., Bouquet, J.-M., Danks, G., Poulain, J., Campsteijn, C., Adamski, M., Cross, I., Yadetie, F., Muffato, M., Louis, A., Butcher, S., Tsagkogeorga, G., Konrad, A., Singh, S., Jensen, M. F., Cong, E. H., Eikeseth-Otteraa, H., Noel, B., Anthouard, V., Porcel, B. M., Kachouri-Lafond, R., Nishino, A., Ugolini, M., Chourrout, P., Nishida, H., Aasland, R., Huzurbazar, S., Westhof, E., Delsuc, F., Lehrach, H., Reinhardt, R., Weissenbach, J., Roy, S. W., Artiguenave, F., Postlethwait, J. H., Manak, J. R., Thompson, E. M., Jaillon, O., Du Pasquier, L., Boudinot, P., Liberles, D. a., Volff, J.-N., Philippe, H., Lenhard, B., Crollius, H. R., Wincker, P., and Chourrout, D. (2010). Plasticity of Animal Genome Architecture Unmasked by Rapid Evolution of a Pelagic Tunicate. Science, 1381(2010).

Dunn, C. W., Hejnol, A., Matus, D. Q., Pang, K., Browne, W. E., Smith, S. a., Seaver, E., Rouse, G. W., Obst, M., Edgecombe, G. D., Sørensen, M. V., Haddock, S. H. D., Schmidt-Rhaesa, A., Okusu, A., Kristensen, R. M., Wheeler, W. C., Martindale, M. Q., and Giribet, G. (2008). Broad phylogenomic sampling improves resolution of the animal tree of life. Nature, 452(7188):745–9.

Dunn, C. W., Leys, S. P., and Haddock, S. H. D. (2015). The hidden biology of sponges and ctenophores. Trends in Ecology & Evolution, pages 1–10.

Ebersberger, I., Strauss, S., and von Haeseler, A. (2009). HaMStR: profile hidden markov model based search for orthologs in ESTs. BMC evolutionary biology, 9: 157.

Eddy, S. R. (2011). Accelerated profile HMM searches. PLoS Computational Biology, 7(10).

Elliott, G. R. D. and Leys, S. P. (2007). Coordinated contractions effectively expel water from the aquiferous system of a freshwater sponge. Journal of Experimental Biology, 210(21): 3736–3748.

Elliott, G. R. D. and Leys, S. P. (2010). Evidence for glutamate, GABA and NO in coordinating behaviour in the sponge, Ephydatia muelleri (Demospongiae, Spongillidae). The Journal of experimental biology, 213: 2310–2321.

Ellwanger, K., Eich, A., and Nickel, M. (2007). GABA and glutamate specifically induce contractions in the sponge Tethya wilhelma. Journal of Comparative Physiology A: Neuroethology, Sensory, Neural, and Behavioral Physiology, 193(1):1–11.

Ellwanger, K. and Nickel, M. (2006). Neuroactive substances specifically modulate rhythmic body contractions in the nerveless metazoon Tethya wilhelma (Demospongiae, Porifera). Frontiers in zoology, 3: 7.

Fairclough, S. R., Chen, Z., Kramer, E., Zeng, Q., Young, S., Robertson, H.M., Begovic, E., Richter, D. J., Russ, C., Westbrook, M. J., Manning, G., Lang, B. F., Haas, B., Nusbaum, C., and King, N. (2013). Premetazoan genome evolution and the regulation of cell differentiation in the choanoflagellate Salpingoeca rosetta. Genome biology, 14(2):R15.

Fernandez-Valverde, S. L., Calcino, A. D., and Degnan, B. M. (2015). Deep developmental transcriptome sequencing uncovers numerous new genes and enhances gene annotation in the sponge Amphimedon queenslandica. BMC Genomics, 16(1):1–11.

Finn, R. D., Coggill, P., Eberhardt, R. Y., Eddy, S. R., Mistry, J., Mitchell, A. L., Potter, S. C., Punta, M., Qureshi, M., Sangrador-Vegas, A., Salazar, G. A., Tate, J., and Bateman, A. (2016). The Pfam protein families database: towards a more sustainable future. Nucleic Acids Research, 44(D1):D279–D285.

Fortunato, S. a. V., Adamski, M., Ramos, O. M., Leininger, S., Liu, J., Ferrier, D. E. K., and Adamska, M. (2014). Calcisponges have a ParaHox gene and dynamic expression of dispersed NK homeobox genes. Nature, 514(7524):620–623.

Ge, J., Li, W., Zhao, Q., Li, N., Chen, M., Zhi, P., Li, R., Gao, N., Xiao, B., and Yang, M. (2015). Architecture of the mammalian mechanosensitive Piezo1 channel. Nature, 527(5): 64–69.

Geng, Y., Bush, M., Mosyak, L., Wang, F., and Fan, Q. R. (2013). Structural mechanism of ligand activation in human GABA(B) receptor. Nature, 504(7479):254–9.

Grabherr, M. G., Haas, B. J., Yassour, M., Levin, J. Z., Thompson, D. a., Amit, I., Adiconis, X., Fan, L., Raychowdhury, R., Zeng, Q., Chen, Z., Mauceli, E., Hacohen, N., Gnirke, A., Rhind, N., di Palma, F., Birren, B. W., Nusbaum, C., Lindblad-Toh, K., Friedman, N., and Regev, A. (2011). Full-length transcriptome assembly from RNA-Seq data without a reference genome. Nature biotechnology, 29(7): 644–52.

Grell, K. G. and Benwitz, G. (1974). Spezifische Verbindungsstrukuren der Faserzellen von Trichoplax adhaerens F.E. Schulze. Z. Naturforsch., 29c:790.

Gur Barzilai, M., Reitzel, A. M., Kraus, J. E. M., Gordon, D., Technau, U., Gurevitz, M., and Moran, Y. (2012). Convergent Evolution of Sodium Ion Selectivity in Metazoan Neuronal Signaling. Cell Reports, 2(2): 242–248.

Haas, B. J., Papanicolaou, A., Yassour, M., Grabherr, M., Blood, P. D., Bowden, J., Couger, M. B., Eccles, D., Li, B., Lieber, M., Macmanes, M. D., Ott, M., Orvis, J., Pochet, N., Strozzi, F., Weeks, N., Westerman, R., William, T., Dewey, C. N., Henschel, R., Leduc, R. D., Friedman, N., and Regev, A. (2013). De novo transcript sequence reconstruction from RNA-seq using the Trinity platform for reference generation and analysis. Nature protocols, 8(8): 1494–512.

Harbison, G. R. (1985). On the classification and evolution of the Ctenophora. In Conway Morris, S. C., George, J. D., Gibson, R., and Platt, H. M., editors, The Origin and Relationships of Lower Invertebrates, pages 78–100. Clarendon Press, Oxford.

Hoff, K. J. and Stanke, M. (2013). WebAUGUSTUS–a web service for training AUGUSTUS and predicting genes in eukaryotes. Nucleic Acids Research, 41(W1):W123–W128.

Huang, S., Chen, Z., Huang, G., Yu, T., Yang, P., Li, J., Fu, Y., Yuan, S., Chen, S., and Xu, A. (2012). HaploMerger: Reconstructing allelic relationships for polymorphic diploid genome assemblies. Genome Research, 22(8): 1581–1588.

Jarvis, E. D., Mirarab, S., Aberer, A. J., Li, B., Houde, P., Li, C., Ho, S. Y. W., Faircloth, B. C., Nabholz, B., Howard, J. T., Suh, A., Weber, C. C., da Fonseca, R. R., Li, J., Zhang, F., Li, H., Zhou, L., Narula, N., Liu, L., Ganapathy, G., Boussau, B., Bayzid, M. S., Zavidovych, V., Subramanian, S., Gabaldon, T., Capella-Gutierrez, S., Huerta-Cepas, J., Rekepalli, B., Munch, K., Schierup, M., Lindow, B., Warren, W. C., Ray, D., Green, R. E., Bruford, M. W., Zhan, X., Dixon, A., Li, S., Li, N., Huang, Y., Derryberry, E. P., Bertelsen, M. F., Sheldon, F. H., Brumfield, R. T., Mello, C. V., Lovell, P. V., Wirthlin, M., Schneider, M. P. C., Prosdocimi, F., Samaniego, J. A., Velazquez, A. M. V., Alfaro-Nunez, A., Campos, P. F., Petersen, B., Sicheritz-Ponten, T., Pas, A., Bailey, T., Scofield, P., Bunce, M., Lambert, D. M., Zhou, Q., Perelman, P., Driskell, A. C., Shapiro, B., Xiong, Z., Zeng, Y., Liu, S., Li, Z., Liu, B., Wu, K., Xiao, J., Yinqi, X., Zheng, Q., Zhang, Y., Yang, H., Wang, J., Smeds, L., Rheindt, F. E., Braun, M., Fjeldsa, J., Orlando, L., Barker, F. K., Jonsson, K. A., Johnson, W., Koepfli, K.-P., O’Brien, S., Haussler, D., Ryder, O. A., Rahbek, C., Willerslev, E., Graves, G. R., Glenn, T. C., McCormack, J., Burt, D., Ellegren, H., Alstrom, P., Edwards, S. V., Stamatakis, A., Mindell, D. P., Cracraft, J., Braun, E. L., Warnow, T., Jun, W., Gilbert, M. T. P., and Zhang, G. (2014). Whole-genome analyses resolve early branches in the tree of life of modern birds. Science, 346(6215):1320–1331.

Jékely, G., Paps, J., and Nielsen, C. (2015). The phylogenetic position of ctenophores and the origin(s) of nervous systems. EvoDevo, 6(1):1.

Katoh, K. and Standley, D. M. (2013). MAFFT multiple sequence alignment software version 7:improvements in performance and usability. Molecular biology and evolution, 30(4): 772–80.

Kim, D., Pertea, G., Trapnell, C., Pimentel, H., Kelley, R., and Salzberg, S. L. (2013). TopHat2: accurate alignment of transcriptomes in the presence of insertions, deletions and gene fusions. Genome biology, 14(4):R36.

King, N., Westbrook, M. J., Young, S. L., Kuo, A., Abedin, M., Chapman, J., Fairclough, S., Hellsten, U., Isogai, Y., Letunic, I., Marr, M., Pincus, D., Putnam, N., Rokas, A., Wright, K. J., Zuzow, R., Dirks, W., Good, M., Goodstein, D., Lemons, D., Li, W., Lyons, J. B., Morris, A., Nichols, S., Richter, D. J., Salamov, A., Sequencing, J. G. I., Bork, P., Lim, W. a., Manning, G., Miller, W. T., McGinnis, W., Shapiro, H., Tjian, R., Grigoriev, I. V., and Rokhsar, D. (2008). The genome of the choanoflagellate Monosiga brevicollis and the origin of metazoans. Nature, 451(7180):783–8.

Krishnan, A., Dnyansagar, R., Almén, M. S., Williams, M. J., Fredriksson, R., Manoj, N., and Schiöth, H. B. (2014). The GPCR repertoire in the demosponge Amphimedon queenslandica: insights into the GPCR system at the early divergence of animals. BMC Evolutionary Biology, 14(1):270.

Krishnan, A. and Schiöth, H. B. (2015). The role of G protein-coupled receptors in the early evolution of neurotransmission and the nervous system. The Journal of experimental biology, 218(Pt 4):562–571.

Langmead, B. and Salzberg, S. L. (2012). Fast gapped-read alignment with Bowtie 2.

Leys, S. P. (2015). Elements of a ‘nervous system’ in sponges. The Journal of experimental biology, 218(Pt 4):581–91.

Leys, S. P., Mackie, G. O., and Meech, R. W. (1999). Impulse conduction in a sponge. The Journal of experimental biology, 202 (Pt 9)(June 1997):1139–1150.

Leys, S. P., Mackie, G. O., and Reiswig, H. M. (2007). The Biology of Glass Sponges. Advances in Marine Biology, 52(06): 1–145.

Li, X., Liu, H., Chu Luo, J., Rhodes, S. a., Trigg, L. M., van Rossum, D. B., Anishkin, A., Diatta, F. H., Sassic, J. K., Simmons, D. K., Kamel, B., Medina, M., Martindale, M. Q., and Jegla, T. (2015). Major diversification of voltage-gated K\n +\n channels occurred in ancestral parahoxozoans. Proceedings of the National Academy of Sciences, page 201422941.

Liebeskind, B. J., Hillis, D. M., and Zakon, H. H. (2011). Evolution of sodium channels predates the origin of nervous systems in animals. Proceedings of the National Academy of Sciences of the United States of America, 108(22): 9154–9159.

Ludeman, D. A., Farrar, N., Riesgo, A., Paps, J., and Leys, S. P. (2014). Evolutionary origins of sensation in metazoans: functional evidence for a new sensory organ in sponges. BMC Evolutionary Biology, 14(3):1–11.

Luo, R., Liu, B., Xie, Y., Li, Z., Huang, W., Yuan, J., He, G., Chen, Y., Pan, Q., Liu, Y., Tang, J., Wu, G., Zhang, H., Shi, Y., Liu, Y., Yu, C., Wang, B., Lu, Y., Han, C., Cheung, D. W., Yiu, S.-M., Peng, S., Xiaoqian, Z., Liu, G., Liao, X., Li, Y., Yang, H., Wang, J., Lam, T.-W., and Wang, J. (2012). SOAPdenovo2: an empirically improved memory-efficient short-read de novo assembler. GigaScience, 1(1):18.

Marçais, G. and Kingsford, C. (2011). A fast, lock-free approach for efficient parallel counting of occurrences of k-mers. Bioinformatics, 27(6): 764–770.

Moran, Y., Barzilai, M. G., Liebeskind, B. J., and Zakon, H. H. (2015). Evolution of voltage-gated ion channels at the emergence of Metazoa. Journal of Experimental Biology, 218: 515–525.

Moran, Y., Fredman, D., Praher, D., Li, X. Z., Wee, L. M., Rentzsch, F., Zamore, P. D., Technau, U., and Seitz, H. (2014). Cnidarian microRNAs frequently regulate targets by cleavage. Genome Research, 24(4): 651–663.

Moroz, L. L. (2015). Convergent evolution of neural systems in ctenophores. Journal of Experimental Biology, 218: 598–611.

Moroz, L. L., Kocot, K. M., Citarella, M. R., Dosung, S., Norekian, T. P., Povolotskaya, I. S., Grigorenko, A. P., Dailey, C., Berezikov, E., Buckley, K. M., Ptitsyn, A., Reshetov, D., Mukherjee, K., Moroz, T. P., Bobkova, Y., Yu, F., Kapitonov, V. V., Jurka, J., Bobkov, Y. V., Swore, J. J., Girardo, D. O., Fodor, A., Gusev, F., Sanford, R., Bruders, R., Kittler, E., Mills, C. E., Rast, J. P., Derelle, R., Solovyev, V. V., Kondrashov, F. a., Swalla, B. J., Sweedler, J. V., Rogaev, E. I., Halanych, K. M., and Kohn, A. B. (2014). The ctenophore genome and the evolutionary origins of neural systems. Nature, 17: 1–123.

Nickel, M. (2010). Evolutionary emergence of synaptic nervous systems: what can we learn from the non-synaptic, nerveless Porifera? Invertebrate Biology, 129(1): 1–16.

Nosenko, T., Schreiber, F., Adamska, M., Adamski, M., Eitel, M., Hammel, J., Maldonado, M., Müller, W. E. G., Nickel, M., Schierwater, B., Vacelet, J., Wiens, M., and Wörheide, G. (2013). Deep metazoan phylogeny: when different genes tell different stories. Molecular phylogenetics and evolution, 67(1): 223–33.

Pérez-Porro, a. R., Navarro-Gómez, D., Uriz, M. J., and Giribet, G. (2013). A NGS approach to the encrusting Mediterranean sponge Crella elegans (Porifera, Demospongiae, Poecilosclerida): transcriptome sequencing, characterization and overview of the gene expression along three life cycle stages. Molecular ecology resources, 454: 494–509.

Pertea, M., Pertea, G. M., Antonescu, C. M., Chang, T.-C., Mendell, J. T., and Salzberg, S. L. (2015). StringTie enables improved reconstruction of a transcriptome from RNA-seq reads. Nature Biotechnology, 33(3).

Petersen, T. N., Brunak, S., von Heijne, G., and Nielsen, H. (2011). SignalP 4.0: discriminating signal peptides from transmembrane regions. Nature methods, 8(10): 785–6.

Philippe, H., Brinkmann, H., Lavrov, D. V., Littlewood, D. T. J., Manuel, M., Wörheide, G., and Baurain, D. (2011). Resolving difficult phylogenetic questions: why more sequences are not enough. PLoS biology, 9(3):e1000602.

Philippe, H., Derelle, R., Lopez, P., Pick, K., Borchiellini, C., Boury-Esnault, N., Vacelet, J., Renard, E., Houliston, E., Quéinnec, E., Da Silva, C., Wincker, P., Le Guyader, H., Leys, S., Jackson, D. J., Schreiber, F., Erpenbeck, D., Morgenstern, B., Wörheide, G., and Manuel, M. (2009). Phylogenomics revives traditional views on deep animal relationships. Current biology: CB, 19(8): 706–12.

Pick, K. S., Philippe, H., Schreiber, F., Erpenbeck, D., Jackson, D. J., Wrede, P., Wiens, M., Alié, A., Morgenstern, B., Manuel, M., and Wörheide, G. (2010). Improved phylogenomic taxon sampling noticeably affects nonbilaterian relationships. Molecular Biology and Evolution, 27(9): 1983–1987.

Pisani, D., Pett, W., Dohrmann, M., Feuda, R., Rota-Stabelli, O., Philippe, H., Lartillot, N., and Wörheide, G. (2015). Genomic data do not support comb jellies as the sister group to all other animals. Proceedings of the National Academy of Sciences, 112(50):201518127.

Ponce, D., Brinkman, D. L., Potriquet, J., and Mulvenna, J. (2016). Tentacle transcriptome and venom proteome of the pacific sea nettle, Chrysaora fuscescens (Cnidaria: Scyphozoa). Toxins, 8(4).

Pratlong, M., Haguenauer, A., Chabrol, O., Klopp, C., Pontarotti, P., and Aurelle, D. (2015). The red coral (Corallium rubrum) transcriptome: a new resource for population genetics and local adaptation studies. Molecular Ecology Resources, 15(5): 1205–1215.

Price, M. N., Dehal, P. S., and Arkin, A. P. (2010). FastTree 2 - Approximately maximum-likelihood trees for large alignments. PLoS ONE, 5(3).

Putnam, N. H., Butts, T., Ferrier, D. E. K., Furlong, R. F., Hellsten, U., Kawashima, T., Robinson-Rechavi, M., Shoguchi, E., Terry, A., Yu, J.-K., Benito-Gutiérrez, E. L., Dubchak, I., Garcia-Fernàndez, J., Gibson-Brown, J. J., Grigoriev, I. V., Horton, A. C., de Jong, P. J., Jurka, J., Kapitonov, V. V., Kohara, Y., Kuroki, Y., Lindquist, E., Lucas, S., Osoegawa, K., Pennacchio, L. a., Salamov, A. a., Satou, Y., Sauka-Spengler, T., Schmutz, J., Shin-I, T., Toyoda, A., Bronner-Fraser, M., Fujiyama, A., Holland, L. Z., Holland, P. W. H., Satoh, N., and Rokhsar, D. S. (2008). The amphioxus genome and the evolution of the chordate karyotype. Nature, 453(7198):1064–71.

Qiu, F., Ding, S., Ou, H., Wang, D., Chen, J., and Miyamoto, M. M. (2015). Transcriptome changes during the life cycle of the red sponge, Mycale phyllophila (Porifera, Demospongiae, Poecilosclerida). Genes, 6(4): 1023–1052.

Ramoino, P., Ledda, F. D., Ferrando, S., Gallus, L., Bianchini, P., Diaspro, A., Fato, M., Tagliafierro, G., and Manconi, R. (2010). Metabotropic??-aminobutyric acid (GABAB) receptors modulate feeding behavior in the calcisponge Leucandra aspera. Journal of Experimental Zoology Part A: Ecological Genetics and Physiology, 313A(3):132–140.

Riesgo, A., Farrar, N., Windsor, P. J., Giribet, G., and Leys, S. P. (2014a). The Analysis of Eight Transcriptomes from All Poriferan Classes Reveals Surprising Genetic Complexity in Sponges. Molecular biology and evolution.

Riesgo, A., Peterson, K., Richardson, C., Heist, T., Strehlow, B., McCauley, M., Cotman, C., Hill, M., and Hill, A. (2014b). Transcriptomic analysis of differential host gene expression upon uptake of symbionts: a case study with Symbiodinium and the major bioeroding sponge Cliona varians. BMC genomics, 15(1):376.

Ryan, J. F., Pang, K., Mullikin, J. C., Martindale, M. Q., and Baxevanis, A. D. (2010). The homeodomain complement of the ctenophore Mnemiopsis leidyi suggests that Ctenophora and Porifera diverged prior to the ParaHoxozoa. EvoDevo, 1(1):9.

Ryan, J. F., Pang, K., Schnitzler, C. E., a. D. Nguyen, A.-d., Moreland, R. T., Simmons, D. K., Koch, B. J., Francis, W. R., Havlak, P., Smith, S. a., Putnam, N. H., Haddock, S. H. D., Dunn, C. W., Wolfsberg, T. G., Mullikin, J. C., Martindale, M. Q., Baxevanis, A. D., Comparative, N., and Program, S. (2013). The Genome of the Ctenophore Mnemiopsis leidyi and Its Implications for Cell Type Evolution. Science, 342(6164):1242592–1242592.

Ryu, T., Seridi, L., Moitinho-Silva, L., Oates, M., Liew, Y. J., Mavromatis, C., Wang, X., Haywood, A., Lafi, F. F., Kupresanin, M., Sougrat, R., Alzahrani, M. A., Giles, E., Ghosheh, Y., Schunter, C., Baumgarten, S., Berumen, M. L., Gao, X., Aranda, M., Foret, S., Gough, J., Voolstra, C. R., Hentschel, U., and Ravasi, T. (2016). Hologenome analysis of two marine sponges with different microbiomes. BMC genomics, 17(1):158.

Sáez, A. G., Lozano, E., and Zaldívar-Riverón, A. (2009). Evolutionary history of Na,K-ATPases and their osmoregulatory role. Genetica, 136(3): 479–490.

Sahlin, K., Vezzi, F., Nystedt, B., Lundeberg, J., and Arvestad, L. (2014). BESST - Efficient scaffolding of large fragmented assemblies. BMC Bioinformatics, 15(1):281.

Sara, M., Sara, A., Nickel, M., and Brümmer, F. (2001). Three New Species of Tethya (Porifera: Demospongia) from German Aquaria. Stuttgarter Beiträge zur Naturkunde, 631(15S):1–16.

Schierwater, B., Kolokotronis, S., Eitel, M., and DeSalle, R. (2009). The Diploblast-Bilateria Sister hypothesis. Communicative & Integrative Biology, 2(5): 1–3.

Schuler, a., Schmitz, G., Reft, A., Ozbek, S., Thurm, U., and Bornberg-Bauer, E. (2015). The rise and fall of TRP-N, an ancient family of mechanogated ion channels, in Metazoa. Genome Biology and Evolution, 7(6): 1–27.

Simakov, O., Kawashima, T., Marlétaz, F., Jenkins, J., Koyanagi, R., Mitros, T., Hisata, K., Bredeson, J., Shoguchi, E., Gyoja, F., Yue, J.-X., Chen, Y.-C., Freeman, R. M., Sasaki, A., Hikosaka-Katayama, T., Sato, A., Fujie, M., Baughman, K. W., Levine, J., Gonzalez, P., Cameron, C., Fritzenwanker, J. H., Pani, A. M., Goto, H., Kanda, M., Arakaki, N., Yamasaki, S., Qu, J., Cree, A., Ding, Y., Dinh, H. H., Dugan, S., Holder, M., Jhangiani, S. N., Kovar, C. L., Lee, S. L., Lewis, L. R., Morton, D., Nazareth, L. V., Okwuonu, G., Santibanez, J., Chen, R., Richards, S., Muzny, D. M., Gillis, A., Peshkin, L., Wu, M., Humphreys, T., Su, Y.-H., Putnam, N. H., Schmutz, J., Fujiyama, A., Yu, J.-K., Tagawa, K., Worley, K. C., Gibbs, R. A., Kirschner, M. W., Lowe, C. J., Satoh, N., Rokhsar, D. S., and Gerhart, J. (2015). Hemichordate genomes and deuterostome origins. Nature, pages 1–19.

Simakov, O., Marletaz, F., Cho, S.-J., Edsinger-Gonzales, E., Havlak, P., Hellsten, U., Kuo, D.-H., Larsson, T., Lv, J., Arendt, D., Savage, R., Osoegawa, K., de Jong, P., Grimwood, J., Chapman, J. a., Shapiro, H., Aerts, A., Otillar, R. P., Terry, A. Y., Boore, J. L., Grigoriev, I. V., Lindberg, D. R., Seaver, E. C., Weisblat, D. a., Putnam, N. H., and Rokhsar, D. S. (2013). Insights into bilaterian evolution from three spiralian genomes. Nature, 493(7433):526–31.

Simão, F. A., Waterhouse, R. M., Ioannidis, P., and Kriventseva, E. V. (2015). BUSCO: assessing genome assembly and annotation completeness with single-copy orthologs. Genome analysis, 31(June):9–10.

Simion, P., Philippe, H., Baurain, D., Jager, M., Richter, D. J., Di Franco, A., Roure, B., Satoh, N., Quéinnec, É., Ereskovsky, A., Lapébie, P., Corre, E., Delsuc, F., King, N., Wörheide, G., and Manuel, M. (2017). A Large and Consistent Phylogenomic Dataset Supports Sponges as the Sister Group to All Other Animals. Current Biology, pages 1–10.

Smith, S. M. E., Morgan, D., Musset, B., Cherny, V. V., Place, A. R., Hastings, J. W., and DeCoursey, T. E. (2011). Voltage-gated proton channel in a dinoflagellate. Proceedings of the National Academy of Sciences, 108(44): 18162–18167.

Sreedharan, S., Shaik, J. H. A., Olszewski, P. K., Levine, A. S., Schiöth, H. B., and Fredriksson, R. (2010). Glutamate, aspartate and nucleotide transporters in the SLC17 family form four main phylogenetic clusters: evolution and tissue expression. BMC genomics, 11(iii):17.

Srivastava, M., Begovic, E., Chapman, J., Putnam, N. H., Hellsten, U., Kawashima, T., Kuo, A., Mitros, T., Salamov, A., Carpenter, M. L., Signorovitch, A. Y., Moreno, M. a., Kamm, K., Grimwood, J., Schmutz, J., Shapiro, H., Grigoriev, I. V., Buss, L. W., Schierwater, B., Dellaporta, S. L., and Rokhsar, D. S. (2008). The Trichoplax genome and the nature of placozoans. Nature, 454(7207):955–60.

Srivastava, M., Simakov, O., Chapman, J., Fahey, B., Gauthier, M. E. a., Mitros, T., Richards, G. S., Conaco, C., Dacre, M., Hellsten, U., Larroux, C., Putnam, N. H., Stanke, M., Adamska, M., Darling, A., Degnan, S. M., Oakley, T. H., Plachetzki, D. C., Zhai, Y., Adamski, M., Calcino, A., Cummins, S. F., Goodstein, D. M., Harris, C., Jackson, D. J., Leys, S. P., Shu, S., Woodcroft, B. J., Vervoort, M., Kosik, K. S., Manning, G., Degnan, B. M., and Rokhsar, D. S. (2010).The Amphimedon queenslandica genome and the evolution of animal complexity. Nature, 466(7307):720–6.

Stamatakis, A. (2014). RAxML version 8: a tool for phylogenetic analysis and post-analysis of large phylogenies. Bioinformatics, 30(9): 1312–1313.

Stanke, M., Diekhans, M., Baertsch, R., and Haussler, D. (2008). Using native and syntenically mapped cDNA alignments to improve de novo gene finding. Bioinformatics, 24(5):637–644.

Stein, W. D. (1995). The sodium pump in the evolution of animal cells. Philosophical transactions of the Royal Society of London. Series B, Biological sciences, 349(1329):263–9.

Strous, M., Kraft, B., Bisdorf, R., and Tegetmeyer, H. E. (2012). The binning of metagenomic contigs for microbial physiology of mixed cultures. Frontiers in Microbiology, 3(DEC):1–11.

Suga, H., Chen, Z., de Mendoza, A., Sebé-Pedrós, A., Brown, M. W., Kramer, E., Carr, M., Kerner, P., Vervoort, M., Sánchez-Pons, N., Torruella, G., Derelle, R., Manning, G., Lang, B. F., Russ, C., Haas, B. J., Roger, A. J., Nusbaum, C., and Ruiz-Trillo, I. (2013). The Capsaspora genome reveals a complex unicellular prehistory of animals. Nature communications, 4: 2325.

Versluis, D., D’Andrea, M. M., Ramiro Garcia, J., Leimena, M. M., Hugenholtz, F., Zhang, J., Öztürk, B., Nylund, L., Sipkema, D., van Schaik, W., de Vos, W. M., Kleerebezem, M., Smidt, H., and van Passel, M. W. J. (2015). Mining microbial metatranscriptomes for expression of antibiotic resistance genes under natural conditions. Scientific reports, 5(January):11981.

Whelan, N. V., Kocot, K. M., Moroz, L. L., and Halanych, K. M. (2015). Error, signal, and the placement of Ctenophora sister to all other animals. Proceedings of the National Academy of Sciences, 112(18): 1–6.

Wray, G. A., Smith, A., Peterson, K., Donoghue, P., Benton, M., Thackray, J., Budd, G., Marshall, C., Shu, D., Isozaki, Y., Zhang, X., Han, J., Maruyama, S., Walcott, C., Knoll, A., Walter, M., Narbonne, G., Christie-Blick, N., Raymond, P., Cloud, P., Budd, G., Jensen, S., Briggs, D., Fortey, R., Morris, S. C., Seilacher, A., Bose, P., Pfluger, F., Rasmussen, B., Bengtson, S., Fletcher, I., McNaughton, N., Pecoits, E., Konhauser, K., Aubet, N., Heaman, L., Veroslavsky, G., Stern, R., Gingras, M., Huldtgren, T., Cunningham, J., Yin, C., Stampanoni, M., Marone, F., Donoghue, P., Bengtson, S., Gaucher, C., Poire, D., Bossi, J., Bettucci, L., Beri, A., Fedonkin, M., Simonetta, A., Ivantsov, A., Sprigg, R., Glaessner, M., Gehling, J., Seilacher, A., Retallack, G., Narbonne, G., Seilacher, A., Grazhdankin, D., Legouta, A., Li, C., Chen, J., Hua, T., Maloof, A., Tang, F., Bengtson, S., Wang, Y., Wang, X., Yin, C., Yin, Z., Grant, S., Grotzinger, J., Watters, W., Knoll, A., Fedonkin, M., Vickers-Rich, P., Swalla, B., Trusler, P., Hall, M., Xiao, S., Zhang, Y., Knoll, A., Chen, J.-Y., Oliveri, P., Li, C., Zhou, G., Gao, F., Hagadorn, J., Peterson, K., Davidson, E., Hagadorn, J., Chen, L., Xiao, S., Pang, K., Zhou, C., Yuan, X., Bengtson, S., Budd, G., Cunningham, J., Margoliash, E., Brown, R., Richardson, M., Boulter, D., Ramshaw, J., Jefferies, R., Runnegar, B., Knoll, A., Carroll, S., Wray, G., Levinton, J., Shapiro, L., Aris-Brosou, S., Yang, Z., Bromham, L., Rambaut, A., Fortey, R., Cooper, A., Penny, D., Welch, J., Fontanillas, E., Bromham, L., Wheat, C., Wahlberg, N., Schulte, J., Erwin, D., Laflamme, M., Tweedt, S., Sperling, E., Pisani, D., Peterson, K., Filipski, A., Murillo, O., Freydenzon, A., Tamura, K., Kumar, S., Tamura, K., Battistuzzi, F., Billing-Ross, P., Murillo, O., Filipski, A., Kumar, S., Hug, L., Roger, A., Ho, S., Phillips, M., Sanderson, M., Thorne, J., Kishino, H., Painter, I., Sanderson, M., Britton, T., Anderson, C., Jacquet, D., Lundqvist, S., Bremer, K., Shaul, S., Graur, D., Battistuzzi, F., Billing-Ross, P., Murillo, O., Filipski, A., Kumar, S., Mello, B., Schrago, C., Rambaut, A., Bromham, L., Kishino, H., Thorne, J., Bruno, W., Douzery, E., Snell, E., Bapteste, E., Delsuc, F., Philippe, H., Battistuzzi, F., Filipski, A., Hedges, S., Kumar, S., Schwartz, R., Mueller, R., Ho, S., Phillips, M., Drummond, A., Cooper, A., Blair, J., Hedges, S., Warnock, R., Yang, Z., Donoghue, P., Battistuzzi, F., Billing-Ross, P., Murillo, O., Filipski, A., Kumar, S., Ayala, F., Rzhetsky, A., Ayala, F., Peterson, K., Lyons, J., Nowak, K., Takacs, C., Wargo, M., McPeek, M., Cutler, D., Chernikova, D., Motamedi, S., Csuros, M., Koonin, E., Rogozin, I., Richter, D., King, N., Dunn, C., Giribet, G., Edgecombe, G., Hejnol, A., Emes, R., Grant, S., Burkhardt, P., Hejnol, A., Martindale, M., Balavoine, G., Adoutte, A., Hirth, F., Miller, D., Ball, E., Northcutt, R., Strausfeld, N., Hirth, F., Tosches, M., Arendt, D., Lyons, T., Reinhard, C., Planavsky, N., Planavsky, N., Sperling, E., Frieder, C., Raman, A., Girguis, P., Levin, L., Knoll, A., Stanley, S., Peterson, K., Cotton, J., Gehling, J., Pisani, D., Butterfield, N., Benito-Gutierrez, E., Arendt, D., Holland, L., Carvalho, J., Escriva, H., Laudet, V., Schubert, M., Shimeld, S., Yu, J.-K., Turner, S., Young, J., Pisani, D., Poling, L., Lyons-Weiler, M., Hedges, S., Otsuka, J., Sugaya, N., Hedges, S., Blair, J., Venturi, M., Shoe, J., and Gu, X. (2015). Molecular clocks and the early evolution of metazoan nervous systems. Philosophical transactions of the Royal Society of London. Series B, Biological sciences, 370(1684):424–431.

Wu, T. D. and Watanabe, C. K. (2005). GMAP: a genomic mapping and alignment program for mRNA and EST sequences. Bioinformatics, 21(9): 1859–1875.

Zakon, H. H. (2012). Adaptive evolution of voltage-gated sodium channels: the first 800 million years. Proceedings of the National Academy of Sciences of the United States of America, 109 Suppl(Supplement_1):10619–25.

Zapata, F., Goetz, F. E., Smith, S. A., Howison, M., Siebert, S., Church, S. H., Sanders, S. M., Ames, C. L., McFadden, C. S., France, S. C., Daly, M., Collins, A. G., Haddock, S. H. D., Dunn, C. W., and Cartwright, P. (2015). Phylogenomic Analyses Support Traditional Relationships within Cnidaria. Plos One, 10(10):e0139068.

